# On the discovered Cancer Driving Nucleotides (CDNs)–Distributions across genes, cancer types and patients

**DOI:** 10.1101/2024.05.29.596367

**Authors:** Lingjie Zhang, Tong Deng, Zhongqi Liufu, Xiangnyu Chen, Shijie Wu, Xueyu Liu, Changhao Shi, Bingjie Chen, Zheng Hu, Qichun Cai, Chenli Liu, Mengfeng Li, Miles E. Tracy, Xuemei Lu, Chung-I Wu, Haijun Wen

## Abstract

A central goal of cancer genomics is to identify, in each patient, all the cancer driving mutations. Among them, point mutations are referred to as Cancer Driving Nucleotides (CDNs), which recur in cancers. The companion study shows that the probability of *i* recurrent hits in **n** patients would decrease exponentially with *i*; hence, any mutation with *i* ≥ 3 hits in the TCGA database is a high-probability CDN. This study characterizes the 50∼150 CDNs identifiable for each cancer type of TCGA (while anticipating 10 times more undiscovered ones) as follows: **i**) CDNs tend to code for amino acids of divergent chemical properties. **ii**) At the genic level, far more CDNs (>5-fold) fall on non-canonical than canonical cancer driving genes (CDGs). Most undiscovered CDNs are expected to be on unknown CDGs. **iii**) CDNs tend to be more widely shared among cancer types than canonical CDGs, mainly because of the higher resolution at the nucleotide than the whole-gene level. **iv**) Most important, among the 50∼100 coding region mutations carried by a cancer patient, 5∼8 CDNs are expected but only 0∼2 CDNs have been identified at present. This low level of identification has hampered functional test and gene targeted therapy. We show that, by expanding the sample size to 10^5^, most CDNs can be identified. Full CDN identification will then facilitate the design of patient-specific targeting against multiple CDN-harboring genes.

## INTRODUCTION

Tumorigenesis in each patient is driven by mutations in the patient’s genome. Hence, a central goal of cancer genomics is to identify *all* driving mutations in each patient. This task is particularly challenging because each driving mutation is present in only a small fraction of patients. As the number of driver mutations in each patient has been estimated to be >5 (Armitage and Doll 1954; Bozic et al. 2010; Hanahan and Weinberg 2011; Belikov 2017; Anandakrishnan et al. 2019), the total number of driver mutations summed over all patients must be quite high.

This study, together with the companion paper (Zhang et al. 2024), are based on one simple premise: In the massively repeated evolution of cancers, any advantageous cancer-driving mutation should recur frequently, say *i* times in **n** patients. The converse that non-recurrent mutations are not advantageous is part of the same premise. We focus on point mutations, referred to as Cancer Driving Nucleotides (CDNs), and formulate the maximum of *i* (denoted *i*\*) in **n** patients if mutations are not advantageous. For example, in the TCGA database with **n** generally in the range of 500∼1000, *i*\* = 3. Hence, any point mutation with *i* ≥ 3 is a CDN. At present, a CDN would have a prevalence of 0.3% among cancer patients. If the sample size approaches 10^6^, a CDN only needs to be prevalent at 5×10^-5^, the theoretical limit (Zhang et al. 2024).

Although there are many other driver mutations (e.g., fusion genes, chromosomal aberrations, epigenetic changes, etc.), CDNs should be sufficiently numerous and quantifiable to lead to innovations in functional tests and treatment strategies. Given the current sample sizes of various databases (Cerami et al. 2012; Weinstein et al. 2013; Tate et al. 2019; de Bruijn et al. 2023), each cancer type has yielded 50∼150 CDNs while the CDNs to be discovered should be at least 10 times more numerous. The number of CDNs currently observed in each patient is 0∼2 for most cancer types. This low-level of discovery has limited functional studies and hampered treatment strategies.

While we are proposing the scale-up of sample size to discover most CDNs, we now characterize CDNs that have been discovered. The main issues are the distributions of CDNs among genes, across cancer types and, most important, among patients. In this context, cancer driver genes (CDGs) would be a generic term. We shall use “canonical CDGs” (or conventional CDGs) for the driver genes in the union set of three commonly used lists (Bailey et al. 2018; Sondka et al. 2018; Martínez-Jiménez et al. 2020). In parallel, CDN-harboring genes, referred to “CDN genes”, constitute a new and expanded class of CDGs.

The first issue is that CDNs are not evenly distributed among genes. The canonical cancer drivers such as *TP53*, *KRAS* and *EGFR* tend to have many CDNs. However, the majority of CDNs, especially those yet-to-be-identified ones, may be rather evenly distributed with each gene harboring only 1∼2 CDNs. Hence, the number of genes with tumorigenic potential may be far larger than realized so far. The second issue is the distribution of CDNs and CDGs among cancer types. It is generally understood that the canonical CDGs are not widely shared among cancer types. However, much (but not all) of the presumed cancer-type specificity may be due to low statistical resolution at the genic level.

The third issue concerns the distribution of CDNs among patients. Clearly, the CDN load of a patient is crucial in diagnosis and treatment. However, the conventional diagnosis at the gene level may have two potential problems. One is that many CDNs do not fall in canonical CDGs as signals from one or two CDNs get diluted. Second, a canonical CDG, when mutated, may be mutated at a non-CDN site. In those patients, the said CDG does not drive tumorigenesis. We shall clarify the relationships between CDN mutations and genes that may or may not harbor them.

The characterizations of discovered CDNs are informative and offer a road map for expanding the CDN list. A complete CDN list for each cancer type will be most useful in functional test, diagnosis and treatment. A full list of mutations that drive the evolution of complex traits is at the center of evolutionary genetics. Such phenomena as complex human diseases (e.g., diabetes.) (Vujkovic et al. 2020; Lagou et al. 2023; Xue et al. 2023; Suzuki et al. 2024), the genetics of speciation (Q. Chen et al. 2022; Wang et al. 2022; Wu 2022) and the evolution of viruses in epidemics (Deng et al. 2022; Ruan et al. 2022; Cao et al. 2023; Ruan et al. 2023) are all prime examples in need of a full list). Thanks to their massively repeated evolution, cancers could be the first complex systems well resolved at the genic level.

## RESULTS

In molecular evolution, a gene under positive selection is recognized by its elevated evolutionary rate (**Fig. 1A and 1C**). There have been numerous methods for determining the extent of rate elevation (Li et al. 1985; Nei and Gojobori 1986; Yang and Swanson 2002; Lawrence et al. 2013; Martincorena et al. 2017; Pan et al. 2022; Sherman et al. 2022; Wang et al. 2022; Ruan et al. 2023) and cancer evolution studies have adopted many of them. However, no model has been developed to take advantage of the massively repeated evolution of cancers (**Fig. 1B**), which happens in tens of millions of people at any time.

**Figure 1.**
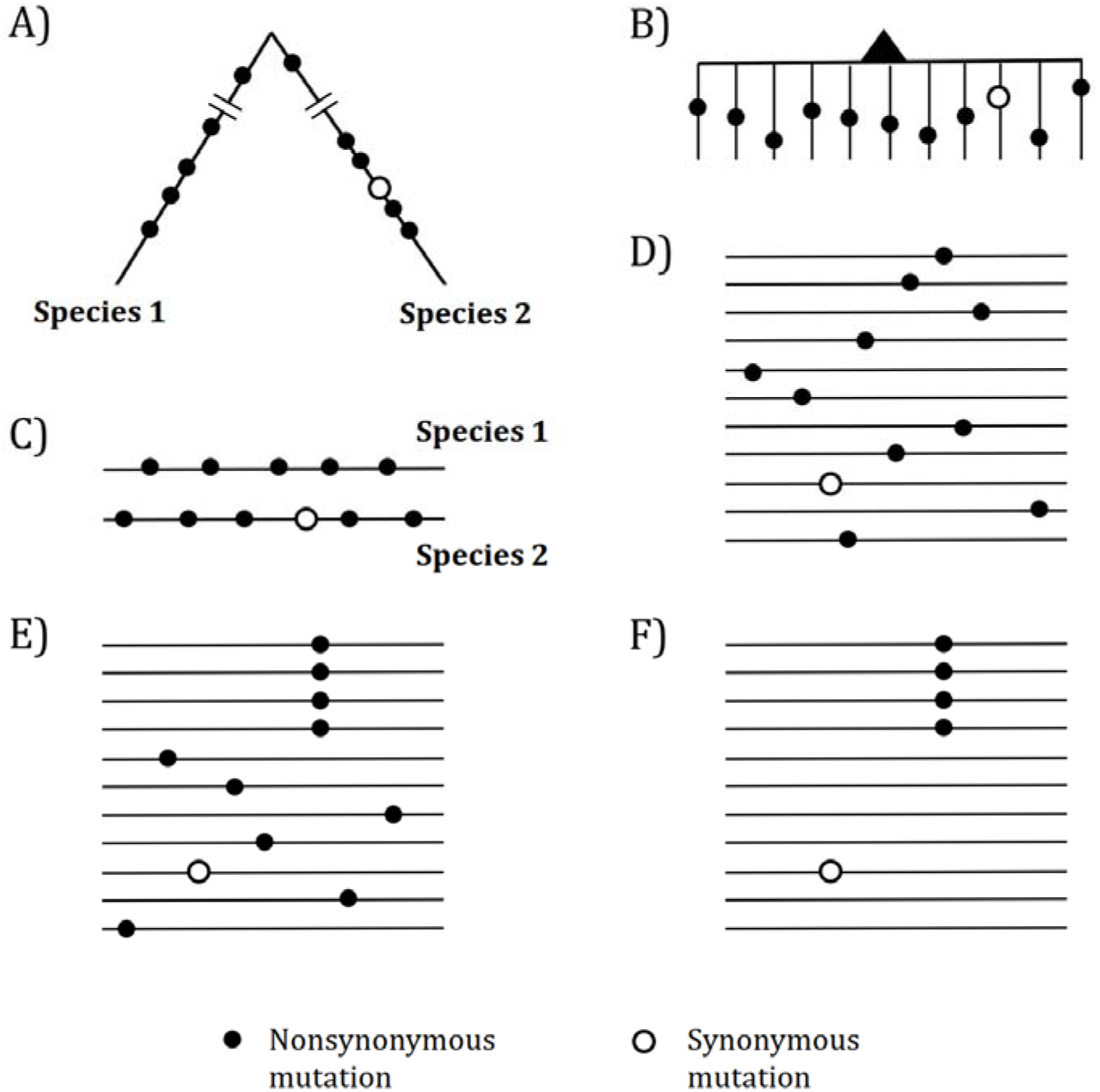
Mutations in organismal evolution vs. cancer evolution. (A, B) A hypothetical example of DNA sequence evolution in organism vs. in cancer with the same number of mutations. (C) Mutation distribution in two species in the organismal evolution of A. (D and E) Mutation distribution in cancer evolution among 10 sequences may have D and E patterns. (F) Another pattern of mutation distribution in cancer evolution with a recurrent site but shows too few total mutations. Mutations of (F) are CDNs missed in the conventional screens.

In the whole-gene analysis, **Fig. 1C-E** are identical, each with *A*: *S* = 10: 1 where *A* and *S* denote nonsynonymous and synonymous mutations, respectively. However, the presence of a 4-hit site in **Fig. 1E** is far less likely to be neutral than **Fig. 1C and 1D**. Although ratio in **Fig. 1F**, *A*: *S* = 4: 1, is statistically indistinguishable from the neutral ratio of about 2.5: 1, **Fig. 1F** in fact has much more power to reject the neutral ratio than **Fig. 1C** and **1D**. After all, the probability that multiple hits are at the same site in a big genome is obviously very small.

## 1. The analyses of CDNs across the whole genome

For the entire coding regions in the cancer genome data, we define ***A***_i_ (or ***S***_i_) as the number of nonsynonymous (or synonymous) sites that harbor a mutation with *i* recurrences. **Table 1** presents the distribution of ***A***_i_ and ***S***_i_ across the 12 cancer types with **n** > 300 (Weinstein et al. 2013).

**Table 1.**
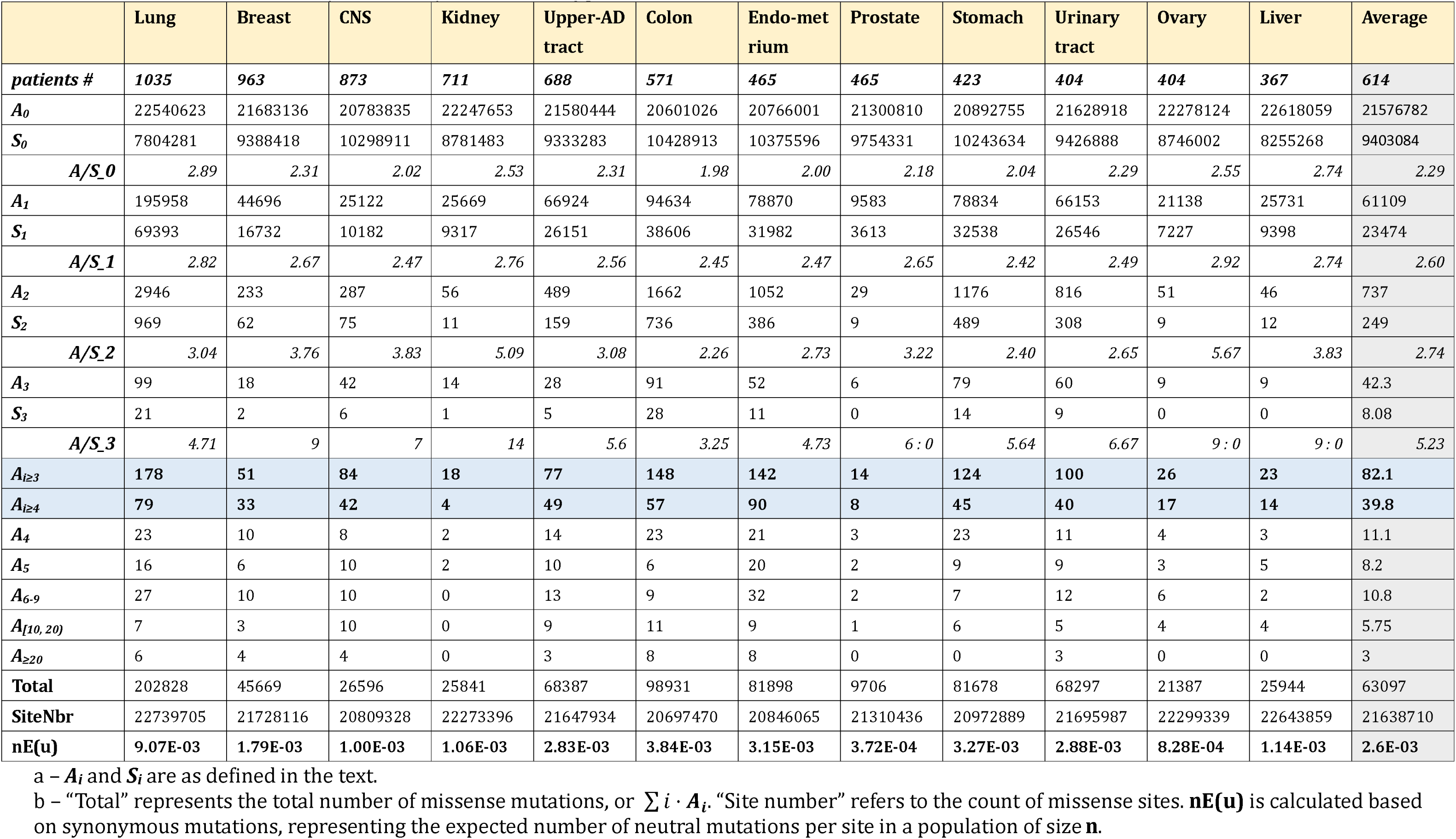
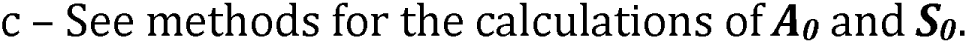
Mutation recurrences (A_i_’s and S_i_’s) in 12 cancer types.

|  | Lung | Breast | CNS | Kidney | Upper-AD tract | Colon | Endo-met rium | Prostate | Stomach | Urinary tract | Ovary | Liver | Average |
| --- | --- | --- | --- | --- | --- | --- | --- | --- | --- | --- | --- | --- | --- |
| **patients #** | **1035** | **963** | **873** | **711** | **688** | **571** | **465** | **465** | **423** | **404** | **404** | **367** | **614** |
| $A_0$ | 22540623 | 21683136 | 20783835 | 22247653 | 21580444 | 20601026 | 20766001 | 21300810 | 20892755 | 21628918 | 22278124 | 22618059 | 21576782 |
| $S_0$ | 7804281 | 9388418 | 10298911 | 8781483 | 9333283 | 10428913 | 10375596 | 9754331 | 10243634 | 9426888 | 8746002 | 8255268 | 9403084 |
| $A/S_0$ | 2.89 | 2.31 | 2.02 | 2.53 | 2.31 | 1.98 | 2.00 | 2.18 | 2.04 | 2.29 | 2.55 | 2.74 | 2.29 |
| $A_1$ | 195958 | 44696 | 25122 | 25669 | 66924 | 94634 | 78870 | 9583 | 78834 | 66153 | 21138 | 25731 | 61109 |
| $S_1$ | 69393 | 16732 | 10182 | 9317 | 26151 | 38606 | 31982 | 3613 | 32538 | 26546 | 7227 | 9398 | 23474 |
| $A/S_1$ | 2.82 | 2.67 | 2.47 | 2.76 | 2.56 | 2.45 | 2.47 | 2.65 | 2.42 | 2.49 | 2.92 | 2.74 | 2.60 |
| $A_2$ | 2946 | 233 | 287 | 56 | 489 | 1662 | 1052 | 29 | 1176 | 816 | 51 | 46 | 737 |
| $S_2$ | 969 | 62 | 75 | 11 | 159 | 736 | 386 | 9 | 489 | 308 | 9 | 12 | 249 |
| $A/S_2$ | 3.04 | 3.76 | 3.83 | 5.09 | 3.08 | 2.26 | 2.73 | 3.22 | 2.40 | 2.65 | 5.67 | 3.83 | 2.74 |
| $A_3$ | 99 | 18 | 42 | 14 | 28 | 91 | 52 | 6 | 79 | 60 | 9 | 9 | 42.3 |
| $S_3$ | 21 | 2 | 6 | 1 | 5 | 28 | 11 | 0 | 14 | 9 | 0 | 0 | 8.08 |
| $A/S_3$ | 4.71 | 9 | 7 | 14 | 5.6 | 3.25 | 4.73 | 6 : 0 | 5.64 | 6.67 | 9 : 0 | 9 : 0 | 5.23 |
| $A_{i \geq 3}$ | 178 | 51 | 84 | 18 | 77 | 148 | 142 | 14 | 124 | 100 | 26 | 23 | 82.1 |
| $A_{i \geq 4}$ | 79 | 33 | 42 | 4 | 49 | 57 | 90 | 8 | 45 | 40 | 17 | 14 | 39.8 |
| $A_4$ | 23 | 10 | 8 | 2 | 14 | 23 | 21 | 3 | 23 | 11 | 4 | 3 | 11.1 |
| $A_5$ | 16 | 6 | 10 | 2 | 10 | 6 | 20 | 2 | 9 | 9 | 3 | 5 | 8.2 |
| $A_{6-9}$ | 27 | 10 | 10 | 0 | 13 | 9 | 32 | 2 | 7 | 12 | 6 | 2 | 10.8 |
| $A_{[10, 20)}$ | 7 | 3 | 10 | 0 | 9 | 11 | 9 | 1 | 6 | 5 | 4 | 4 | 5.75 |
| $A_{\geq 20}$ | 6 | 4 | 4 | 0 | 3 | 8 | 8 | 0 | 0 | 3 | 0 | 0 | 3 |
| **Total** | 202828 | 45669 | 26596 | 25841 | 68387 | 98931 | 81898 | 9706 | 81678 | 68297 | 21387 | 25944 | 63097 |
| **SiteNbr** | 22739705 | 21728116 | 20809328 | 22273396 | 21647934 | 20697470 | 20846065 | 21310436 | 20972889 | 21695987 | 22299339 | 22643859 | 21638710 |
| **nE(u)** | **9.07E-03** | **1.79E-03** | **1.00E-03** | **1.06E-03** | **2.83E-03** | **3.84E-03** | **3.15E-03** | **3.72E-04** | **3.27E-03** | **2.88E-03** | **8.28E-04** | **1.14E-03** | **2.6E-03** |

a – $A_i$ and $S_i$ are as defined in the text.
b – “Total” represents the total number of missense mutations, or $\sum i \cdot A_i$ . “Site number” refers to the count of missense sites. **nE(u)** is calculated based on synonymous mutations, representing the expected number of neutral mutations per site in a population of size **n**.
c – See methods for the calculations of $\mathbf{A}_\theta$ and $\mathbf{S}_\theta$ .

For neutral mutations, we define *i*\* as the threshold above which the expected numbers of ***A***_i_ would be <1, i.e. E[**A**_i≥i*_] 1, The corollary is that all A*_i≥i*_* sites are advantageous CDNs. (Since ***S***_i_ is ∼ ***A***_i_ /2.3, the same *i*\* would apply to ***S***_i_ as well: E[S_i≥i*_] < 1,As *i*\* is a function of the number of patients (**n**), it is shown mathematically in the companion study (Zhang et al. 2024) that *i*\* = 3 for **n** < 1000. Interestingly, while the E[**A***_i≥3_*] is < 1, the expected E[**A***_i≥4_*] is ZZZZ, in the order of 0.001. Hence, *i*^*^ = 4 may be considered unnecessarily stringent.

We should note that this study is constrained by **n** < 1000 in the TCGA databases. (Databases with larger **n**’s are also used where the actual **n**’s are often uncertain.) At *i*\* = 3, we could detect only a fraction (<10%; see below) of CDNs. Many more tumorigenic mutations may be found in the *i* = 1 or 2 classes although not every one of them is a CDN. Since these two classes of mutations are far more numerous, they should account for the bulk of CDNs to be discovered. Indeed, **Table1** shows 76 A*_i≥3_* CDN mutations per cancer type but 681 ***A***_2_ and 56,648 ***A***_1_ mutations in the lower recurrence groups. If **n** reaches 10^5∼6^, most of the undiscovered CDNs in the ***A***_1_ and ***A***_2_ classes should be identified (Zhang et al. 2024).

In Table 2, we estimate the proportion of the ***A***_1_ and ***A***_2_ mutations that are possible CDNs. The relationships of ***A***_3_/***S***_3_ > ***A***_2_/***S***_2_, ***A***_2_/***S***_2_ > ***A***_1_/***S***_1_, and ***A***_1_/***S***_1_ > ***A***_0_/***S***_0_ are almost always observed in **Table 1** with 32 *(3 × 8 + 2 × 4)* out of 36 such relationships. The use of ***A*/*S*** ratios may still under-estimate the selective advantages of ***A***_1∼3_ mutations because ***S***_1∼3_ may have slight advantages as well (Zhang et al. 2024). Assuming ***S***_1_ is truly neutral, we use ***S***_0_ to ***S***_1_ as the basis to calculate the excess of ***A***_1∼3_ in **Table 2** where 35 of the 36 *Obs*(***A***_i_) > *Exp* (***A***_i_) relationships can be observed. The implication is that hundreds and, likely low thousands, of ***A***_1_’s and ***A***_2_’ should be CDNs whereas we have only confidently identified ∼76 strong CDNs, on average, for a cancer type. (Note that ***A***_1_ excesses are less reliable since a 1% error in the calculation would mean 566 CDNs.)

**Table 2.**
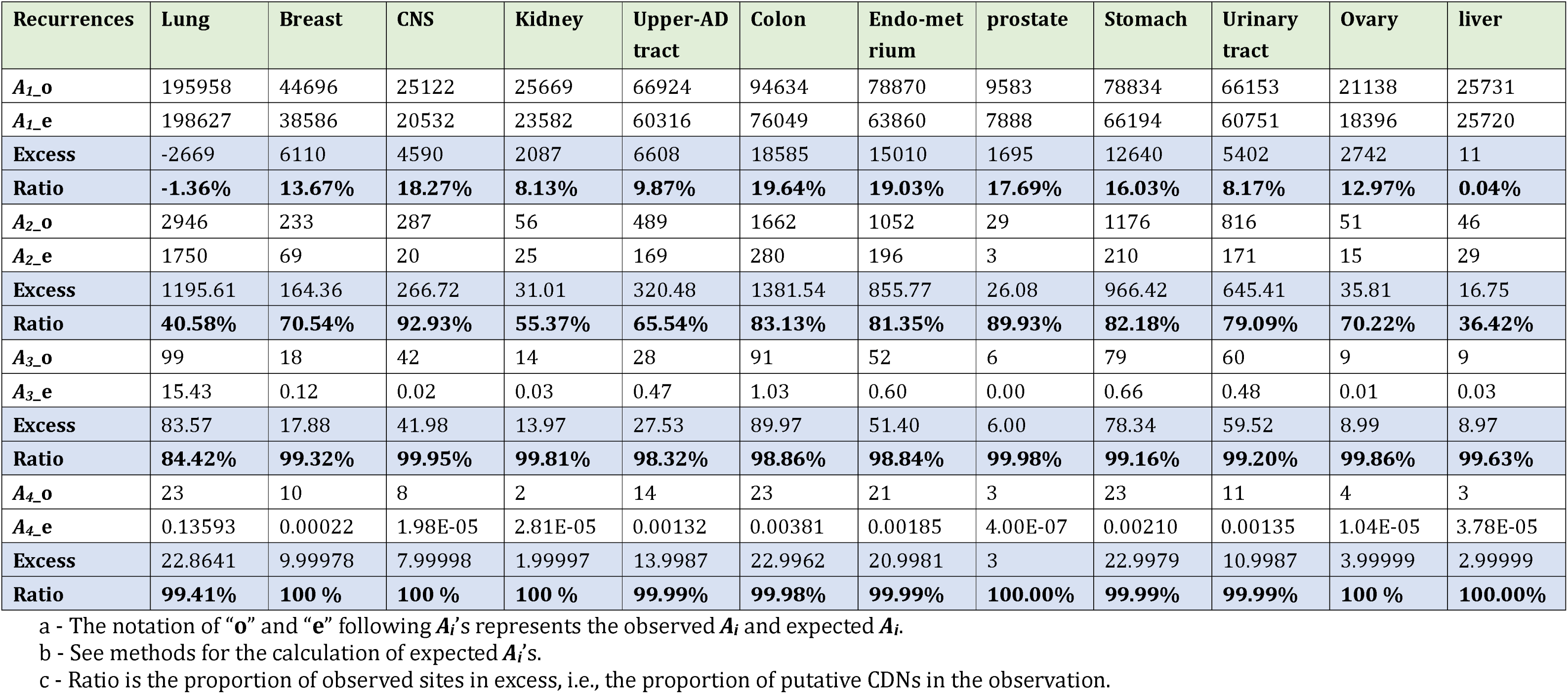
Excess of A_i_’s of each i class.

## 2. CDNs and the amino acids affected

We now ask whether the amino acid changes associated with CDNs bear the signatures of positive selection. Amino acids that have divergent physico-chemical properties have been shown to be under strong selection, both positive and negative (Chen, He, et al. 2019; Chen, Lan, et al. 2019; Q. Chen et al. 2022). We note that, in almost all cases in cancer evolution, when a codon is altered, only one nucleotide of the triplet codon is changed. Among the 190 amino acid (AA, 20×19/2) pairs, only 75 of the pairs differ by one bp (Tang et al. 2004). For example, Pro (CCN) and Ala (GCN) may differ by only one bp but Pro and Gly (GGN) must differ by at least 2 bp. These 75 AA changes, referred to as the elementary AA changes (Grantham 1974; Li et al. 1985; Yang et al. 2003; Meyer et al. 2021), account for almost all AA substitutions in somatic evolution.

In a series of studies (Tang et al. 2004; Chen, He, et al. 2019; Chen, Lan, et al. 2019), we have defined the physico-chemical distances between AAs of the 75 elementary pairs as *ΔUi*, where *i* = 1 to 75. *ΔUi r*eflects 47 measures of AA differences including hydrophobicity, size, charge etc. and ranges between 0 and 1. The most similar pair, Ser and Thr, has *ΔUi*= 0 and the most dissimilar pair is Asp and Try with *ΔUi*= 1. These studies show that *ΔUi*, is a strong determinant of the evolutionary rates of DNA sequences and that large-step changes (i.e., large *ΔUi*,’s) are more acutely “recognized” by natural selection. These large-step changes are either highly deleterious or highly advantageous. Most strikingly, advantageous mutations are enriched with AA pairs of *ΔUi*, > 0.8 (Chen, He, et al. 2019).

To analyze the properties of CDNs, we choose 6 cancer types from **Table 1** that have the largest sample sizes (**n** > 500) but leap over kidney since kidney cancers have unusually low CDN counts. In **Fig. 2**, we divide the CDNs into groups according to the number of recurrences, *i*. CDNs of similar *i*’s are merged into the same group in the descending order of *i*, until there are at least 10 CDNs in the group. The 6 cancer types show two clear trends: 1) the proportion of CDNs with *ΔUi*, > 0.8 (red color segments) increases in groups with higher recurrences; 2) in contrast, the proportion of CDNs with *ΔUi* < 0.4 (green segments) decreases as recurrences increase. These two trends would mean that highly recurrent CDNs tend to involve larger AA distances (*ΔUi* > 0.8) and similar AAs tend not to manifest strong fitness increases. In general, CDNs alter amino acids in ways that expose the changes to strong selection.

**Figure 2.**
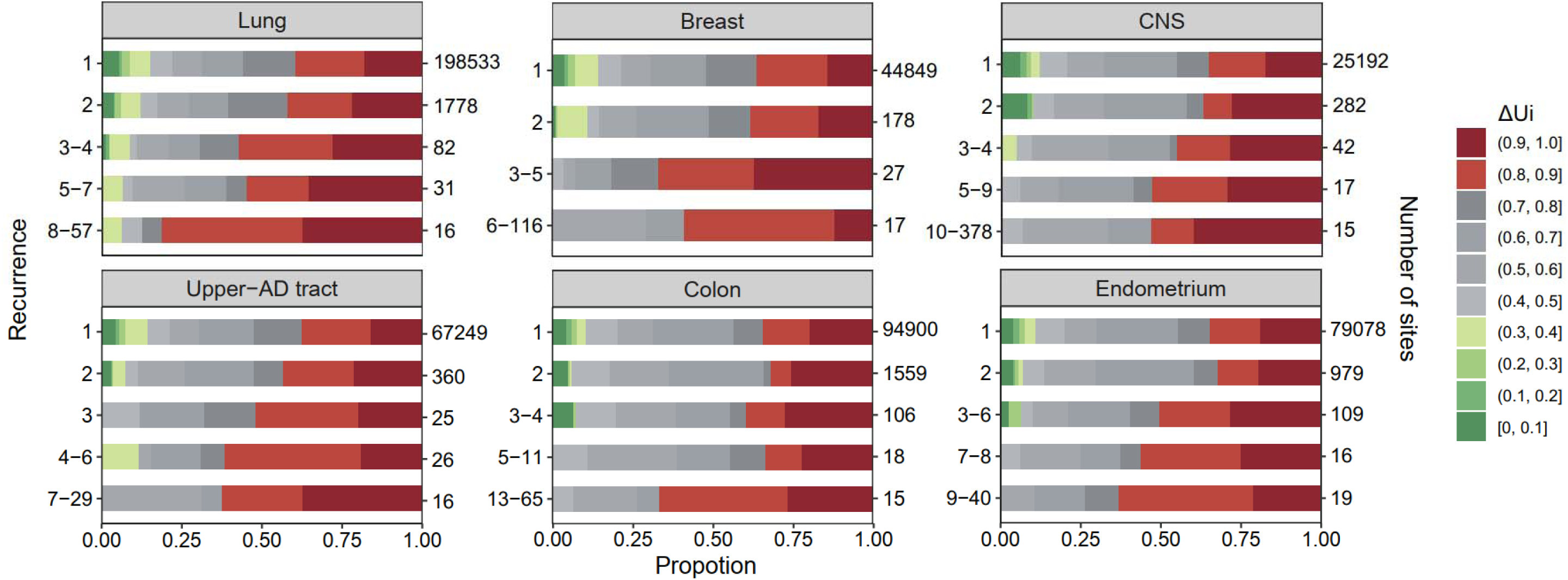
Analysis across 6 cancer types., ranging between 0 and 1 (Tang et al. 2004; Chen, He, et al. 2019), is a measure of p differences among the 20 amino acids (see the text). The most similar amino acids have near 0 and the most dissimilar ones have Each panel corresponds to one cancer type, with horizontal bar represents distribution of each recurrence group. The numbers panel are i values and on the right are the number of sites. Note that the proportion of dark red segments increases as i increases. This that mutations at high recurrence sites (larger i’s) code for amino acids that are chemically very different from the wild type.

## 3. CDNs in relation to the genes harboring them

We shall use the term “CDN genes” for genes having at least one CDN site. Since CDN genes contribute to tumorigenesis when harboring a CDN mutation, they should be considered cancer drivers as well. CDN genes have two desirable qualities for recognition as driver genes. First, CDNs are straightforward and unambiguous to define (e.g., *i* ≥ 3 for **n** < 1000). In the literature, there have been multiple definitions of cancer driver genes (Reimand and Bader 2013; Porta-Pardo and Godzik 2014; Mularoni et al. 2016; Arnedo-Pac et al. 2019), resulting in only modest overlaps among cancer gene lists (see **Fig. S3**). Second, the evolutionary fitness of CDN, and hence the tumorigenic potentials of CDN genes, can be computed (supplement File S1).

We now present the analyses of CDN genes, using the same 6 cancers of **Fig. 2**. Two types of CDN genes are shown in **Table 3**. Type I genes fulfill the conventional criterion of fast evolution with the whole-gene Ka/Ks (or dN/dS) significantly larger than 1 (Martincorena et al. 2017). Averaged across cancer types, Type I overlaps by 95.7% with the canonical CDG list, which is the union of three popular lists (Bailey et al. 2018; Sondka et al. 2018; Martínez-Jiménez et al. 2020). Type I genes are mostly well-known canonical CDGs (e.g., *TP53*, *PIK3CA*, and *EGFR*).

**Table 3.**
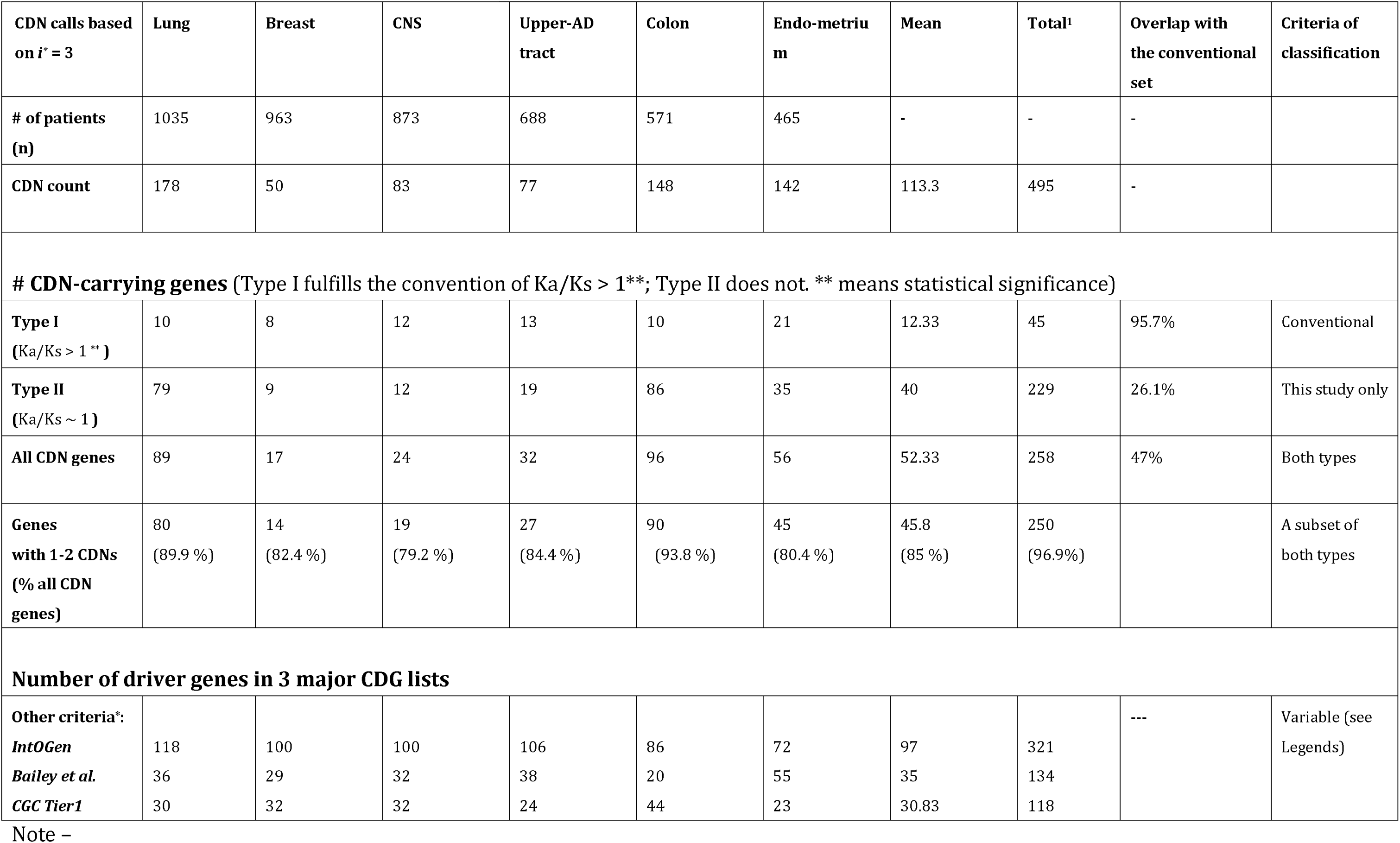
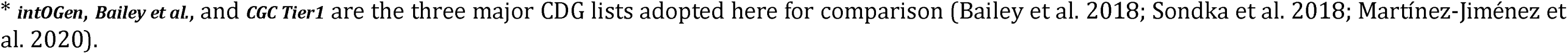
Distribution of CDNs among genes.

| CDN calls based on $i^* = 3$ | Lung | Breast | CNS | Upper-AD tract | Colon | Endo-metrium | Mean | Total<sup>1</sup> | Overlap with the conventional set | Criteria of classification |
| --- | --- | --- | --- | --- | --- | --- | --- | --- | --- | --- |
| # of patients (n) | 1035 | 963 | 873 | 688 | 571 | 465 | - | - | - |  |
| CDN count | 178 | 50 | 83 | 77 | 148 | 142 | 113.3 | 495 | - |  |
| **# CDN-carrying genes** (Type I fulfills the convention of $Ka/Ks > 1^{**}$; Type II does not. **\*\*** means statistical significance) | | | | | | | | | | |
| **Type I**<br>($Ka/Ks > 1^{**}$) | 10 | 8 | 12 | 13 | 10 | 21 | 12.33 | 45 | 95.7% | Conventional |
| **Type II**<br>($Ka/Ks \sim 1$) | 79 | 9 | 12 | 19 | 86 | 35 | 40 | 229 | 26.1% | This study only |
| **All CDN genes** | 89 | 17 | 24 | 32 | 96 | 56 | 52.33 | 258 | 47% | Both types |
| **Genes with 1-2 CDNs**<br>(% all CDN genes) | 80<br>(89.9 %) | 14<br>(82.4 %) | 19<br>(79.2 %) | 27<br>(84.4 %) | 90<br>(93.8 %) | 45<br>(80.4 %) | 45.8<br>(85 %) | 250<br>(96.9%) |  | A subset of both types |
| **Number of driver genes in 3 major CDG lists** |  |  |  |  |  |  |  |  |  |  |
| **Other criteria\*:** |  |  |  |  |  |  |  |  | --- | Variable (see Legends) |
| *IntOGen* | 118 | 100 | 100 | 106 | 86 | 72 | 97 | 321 |  |  |
| *Bailey et al.* | 36 | 29 | 32 | 38 | 20 | 55 | 35 | 134 |  |  |
| *CGC Tier1* | 30 | 32 | 32 | 24 | 44 | 23 | 30.83 | 118 |  |  |

Note –
\* *intOGen*, *Bailey et al.*, and *CGC Tier1* are the three major CDG lists adopted here for comparison (Bailey et al. 2018; Sondka et al. 2018; Martínez-Jiménez et al. 2020).

Type II (CDN genes) is the new class of cancer driver genes. These genes have CDNs but do not meet the conventional criteria of whole-gene analysis. Obviously, if a gene has only one or two CDNs plus some sporadic hits, the whole-gene Ka/Ks would not be significantly greater than 1. As shown in **Table 3**, over 80% of CDN genes have only 1∼2 CDN sites. The salient result is that Type II genes outnumber Type I genes by a ratio of 5: 1 (229: 45, column 8, **Table 3**). Furthermore, Type II genes overlap with the canonical CDG list by only 23%.

Type II genes represent a new class of cancer drivers that concentrate their tumorigenic strength on a small number CDN sites. They have been missed by the conventional whole-gene definition of cancer drivers. One such example is the *FGFR3* gene in lung cancer. This gene of 809 codons has only 8 hits, among which one is a CDN (*i* = 3) in lung cancer. It is noticed solely for this CDN. In the supplemental text, we briefly annotate these new cancer driver genes for comparisons with the canonical driver genes (supplement File S1). Possible functional tests in the future can be found in Discussion.

We now briefly discuss the driver genes listed in previous studies as shown at the lower part of **Table 3** (Bailey et al. 2018; Sondka et al. 2018; Martínez-Jiménez et al. 2020). From the total number of CDGs listed, it is clear that the overlaps are limited. As analyzed before (Wu et al. 2016), conventional gene lists overlap mainly by a core set of high Ka/Ks genes. This core set has not changed much as various criteria such the replication timing, expression profiles, epigenetic features are introduced. These criteria are the reasons for the many CDGs recognized by only a small subset of CDG lists. CDN genes, in contrast, can be objectively defined as CDN mutations (*i* recurrences in **n** samples) themselves are unambiguous.

### Variation in CDN number and tumorigenic contribution among genes

By and large, the distribution of CDNs among genes is very uneven. **Fig. 3A** shows 10 genes with at least 6 CDNs whereas 87 genes have only one CDN. Two genes stand out for the number of CDNs they harbor, *TP53* and *PIK3CA*, which also happen to be the only genes mutated in >15% of all cancer patients surveyed (Kandoth et al. 2013). Clearly, the prevalence of mutations in a gene is a function of the number of strong CDNs it harbors.

**Figure 3.**
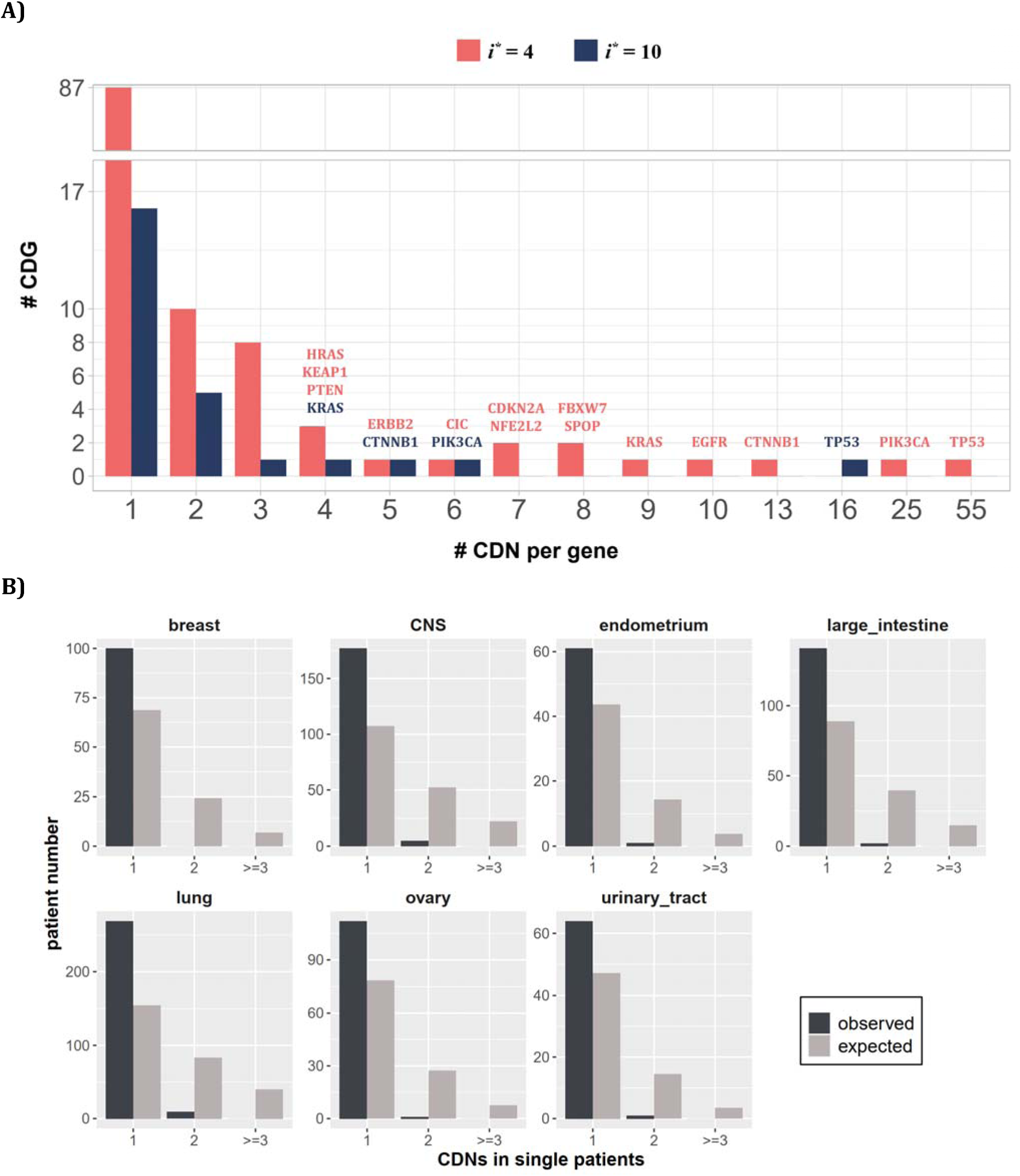
Distribution of CDNs among genes. (A) Out of 119 CDN-carrying genes (red bars), 87 have only one CDN. For the rest, TP53 possesses the most CDNs with three others having more than 10 CDNs. (B) CDN number in TP53 among patients. The dark bar represents the observed patient number with corresponding CDNs of the X-axis. The grey bar shows the expected patient distribution. Clearly, TP53 only needs to contribute one CDN to drive tumorigenesis. Hence, TP53 (and other canonical driver genes; see text), while prevalent, does not contribute disproportionately to the tumorigenesis of each patient.

Although a small number of genes have unusually high number of CDNs, these genes may not drive the tumorigenesis in proportion to their CDN numbers in individual patients. **Fig. 3B** shows the number of CDN mutations on *TP53* that occur in any single patient. Usually, only one CDN change is observed in a patient whereas 2 or 3 CDN mutations are expected. It thus appears that CDNs on the same genes are redundant in their tumorigenic effects such that the second hit may not yield additional advantages. This pattern of disproportionally lower contribution by CDN-rich genes is true in other genes such as *EGFR* and *KRAS*. Consequently, the large number of genes with only 1 or 2 CDN sites are disproportionately important in driving the tumorigenesis of individual patients.

## 4. CDNs in relation to the cancer types - The pan-cancer properties

In the current literature, cancer driver genes (however they are defined) generally meet the statistical criteria for driver genes in only one or a few cancer types. However, genes may in fact contribute to tumorigenesis but are insufficiently prevalent to meet the statistical requirements for CDGs. Many genes are indeed marginally qualified as drivers in some tissues and barely miss the statistical cutoff in others. To see if genes that drive tumorigenesis in multiple tissues are more common than currently understood, we need to raise the sensitivity of cancer driver detection. Thus, CDNs may provide the resolution.

To test the pan-cancer driving capacity of CDNs, we define *i*_max_ as the largest *i* values among the 12 cancer types for each CDN. The number of cancer types where the said mutation can be detected (i.e., *i* > 0) is designated NC12. **Fig. 4** presents the relationship between the observed NC12 of each CDN against *i*_max_ of that CDN. Clearly, many CDNs are observed in multiple cancer types (NC12 > 3), even though they do not qualify as a driver gene in all but a single cancer type. It happens frequently when a mutation has *i* > 3 in one cancer type but has *i* < 3 in others. One extreme example is C394 and G395 in *IDH1*. In CNS, both sites show *i* IZ 3, while in 6 other cancer types (lung, breast, large intestine, prostate, urinary tract, liver), their hits are *i* < 3 but > 0. Conditional on a specific site informed by a cancer type, a mutation in another cancer type should be very unlikely if the mutation is not tumorigenic in multiple tissues. Hence, the pattern in **Fig. 4** is interpreted to be drivers in multiple cancer types, but with varying statistical strength.

**Figure 4.**
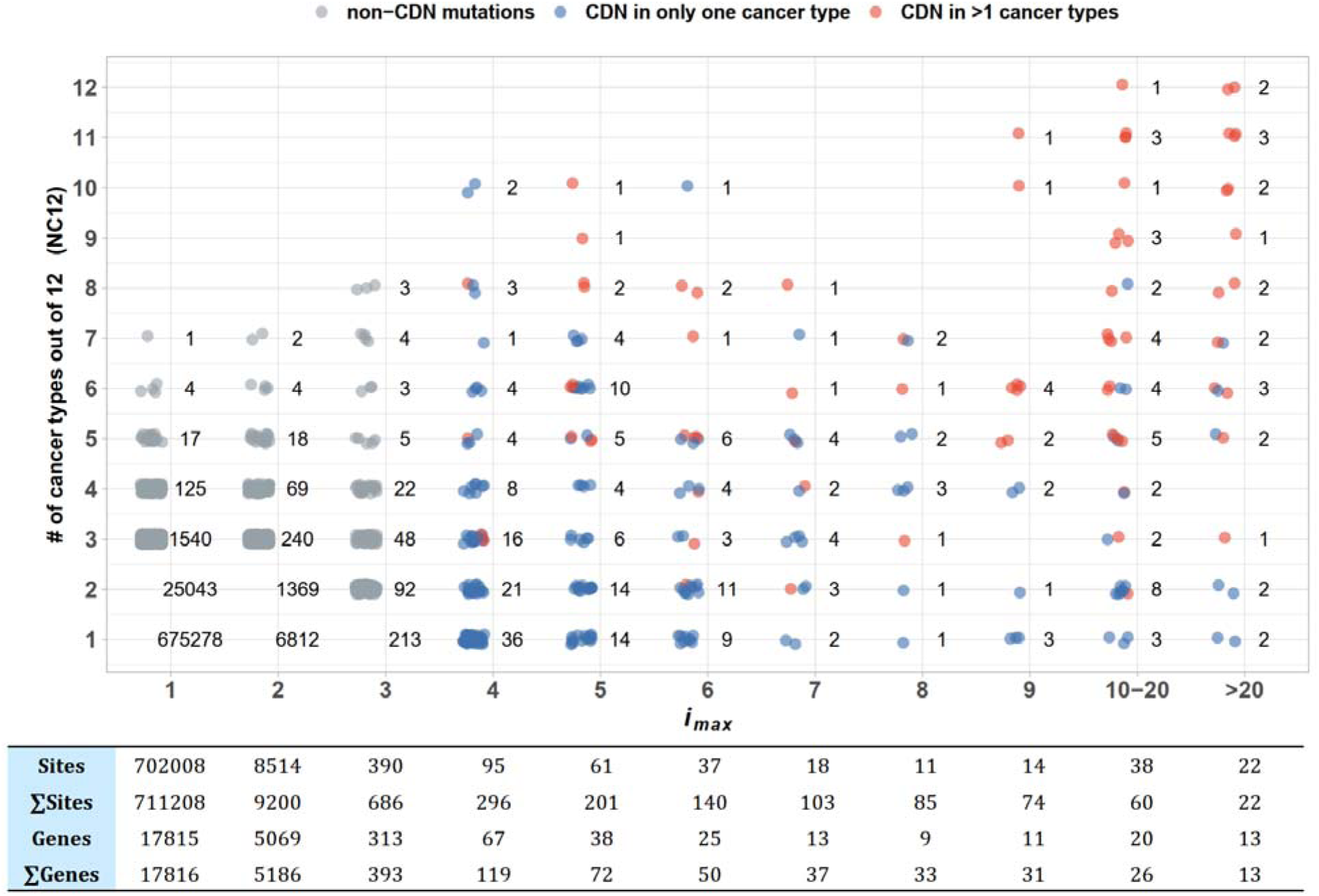
Sharing of CDNs across cancer types. The X-axis shows i_max_, which is the largest i a CDN reaches among the 12 cancer types. The Y axis shows the number of cancer types whre the mutation also occurs. Each dot is a CDN and the number of dots in the cloud is given. The blue and red dots denote, respectively, mutations classified as a CDN in one or multiple cancer types. Grey dots are non-CDNs. The table in the lower panel summarizes the number of sites and the number of genes harboring these sites.

Examining **Fig. 4** more carefully, we could see that CDNs with a larger *i*_max_ in one cancer type are more likely to be identified as CDNs in multiple cancer types (red dots, *r* = 0.97, p = 9.23×10^-5^, Pearson’s correlation test). Of 22 sites with *i*_max_ > 20, 15 are identified as CDNs (*i* ≥ 3) in multiple cancer types, with a median NC12 of 9. On the opposite end, 2 CDNs with *i*_max_ > 20 are observed in only one cancer type (*EGFR*: T2573 in lung and *FGFR2*: C755 in endometrium cancer). The bimodal pattern suggests that a few cancer driver mutations are tissue specific whereas most others appear to have pan-cancer driving potentials.

To conclude, when a driver is observed in more than one cancer type, it is often a cancer driver in many others, but insufficiently powerful to meet the statistical criteria for driver mutations. This pan-cancer property can be seen at the higher resolution of CDN, but is often missed at the whole-gene level. Cancers of the same tissue in different patients, often reported to have divergent mutation profiles (Nik-Zainal et al. 2012; Roberts and Gordenin 2014), should be a good test of this hypothesis.

## 5. CDNs in relation to individual patients and therapeutic strategies

In previous sections, the focus is on the population of cancer patients; for example, how many in the patient population have certain mutations. We now direct the attention to individual patients. It would be necessary to pinpoint the CDN mutations in each patient in order to delineate the specific evolutionary path and to devise the treatment strategy. We shall first address the cancer driving power of CDN vs. non-CDN mutations in the same gene.

*1) Efficacy of targeted therapy against CDNs vs non-CDNs*

In general, a patient would have many point mutations, only a few of which are strong CDNs. We may ask whether most mutations on the canonical genes, such as *EGFR*, are CDNs. Presumably, synonymous, and likely many nonsynonymous, mutations on canonical genes may not be CDNs. It would be logical to hypothesize that patients whose *EGFR* has a CDN mutation (Group1 patients) should benefit from the gene-targeted therapy more than patients with a non-CDN mutation on the same gene (Group2 patients). In the second group, *EFGR* may be a non-driver of tumorigenesis.

Published data (AACR Project GENIE Consortium 2017; Choudhury et al. 2023) are re-analyzed as shown in **Fig. 5**. The hypothesis that patients of Group2 would not benefit as much as those of Group1 is supported by the analysis. This pattern further strengthens the underlying assumption that non-CDN mutations, even on canonical genes, are not cancer drivers.

**Figure 5.**
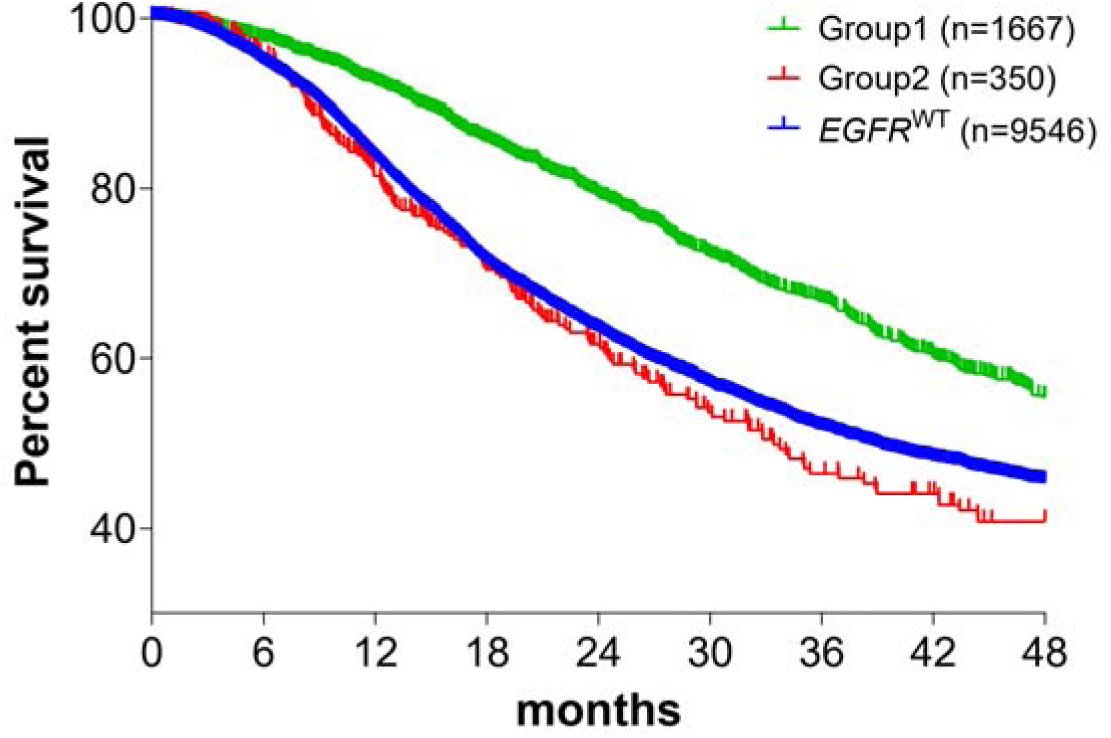
Survival analysis of non-small cell lung cancer (NSCLC) patients based on EGFR mutation status. Patient data were retrieved from the GENIE database (https://genie.cbioportal.org/) and stratified into three groups based on EGFR mutation profiles: Group1 comprises patients with EGFR CDN mutations; Group2 includes patients with nonsynonymous mutations in EGFR that are not CDNs; The EGFR^WT^ group consists of patients with no EGFR mutations (see methods). Patients of Group1 and Group2 received EGFR-targeted therapies in accordance with the guidelines for managing EGFR mutant NSCLC (Passaro et al. 2022; Choudhury et al. 2023). Survival analysis using the Kaplan-Meier method revealed a significantly higher survival rate for Group1 patients compared to Group2 and the EGFR^WT^ group (p < 0.001).

*2) Number of CDNs in each patient*

We postulate that a full set of CDNs should be able to inform about the cause of each cancer as well as the design of gene-targeted therapy. In **Table 4**, the known CDNs based on TCGA are tallied. Note that only a few CDNs fall on the canonical driver genes whereas most CDNs fall on the non-conventional ones.

**Table 4.** Numbers of patients with CDNs vs. number of patients with any non-synonymous mutations in the same genes.

|  | Lung |  | Breast |  | CNS |  | Upper-AD tract |  | Colon |  | Endometrium |  |
| --- | --- | --- | --- | --- | --- | --- | --- | --- | --- | --- | --- | --- |
|  | CDN<sup>1, 3</sup><br>(178) | gene<sup>2, 3</sup><br>(89) | CDN<br>(50) | Gene<br>(17) | CDN<br>(83) | Gene<br>(24) | CDN<br>(77) | Gene<br>(32) | CDN<br>(148) | Gene<br>(96) | CDN<br>(142) | Gene<br>(56) |
| **n<sub>0</sub>** | 342<br>(33%)<sup>4</sup> | 53<br>(5.3%) | 492<br>(51.1%) | 415<br>(43.1%) | 235<br>(26.9%) | 163<br>(18.7%) | 268<br>(39%) | 140<br>(20.3%) | 102<br>(17.9%) | 42<br>(7.4%) | 42<br>(9%) | 14<br>(3%) |
| **n<sub>1</sub>** | 411<br>(39.7%) | 70<br>(6.8%) | 379<br>(39.4%) | 395<br>(41%) | 359<br>(41.1%) | 306<br>(35.1%) | 268<br>(39%) | 229<br>(33.3%) | 159<br>(27.8%) | 79<br>(13.8%) | 108<br>(23.2%) | 59<br>(12.7%) |
| **n<sub>2</sub>** | 192<br>(18.6%) | 84<br>(8.1%) | 73<br>(7.6%) | 114<br>(11.8%) | 225<br>(25.8%) | 293<br>(33.6%) | 101<br>(14.7%) | 171<br>(24.9%) | 140<br>(24.5%) | 93<br>(16.3%) | 169<br>(36.3%) | 101<br>(21.7%) |
| **n<sub>>2</sub>** | 90<br>(8.7%) | 826<br>(79.8%) | 18<br>(1.9%) | 38<br>(3.9%) | 53<br>(6.1%) | 110<br>(12.6%) | 50<br>(7.3%) | 147<br>(21.4%) | 170<br>(29.8%) | 357<br>(62.5%) | 146<br>(31.4%) | 291<br>(62.6%) |
| **Total n** | 1035 | 1035 | 963 | 963 | 873 | 873 | 688 | 688 | 571 | 571 | 465 | 465 |
| **Mean #** | 1.06 | 7.19 | 0.61 | 0.78 | 1.12 | 1.44 | 0.93 | 1.63 | 1.96 | 4.6 | 2.17 | 3.7 |

Note –
<sup>1</sup> - **n<sub>i</sub>** designates the number of patients with *i* CDN mutations.
<sup>2</sup> - In this column, **n<sub>i</sub>** designates the number of patients with any nonsynonymous mutation in the same gene as the CDN column.
<sup>3</sup> - The number in the parentheses is the total number of CDNs or genes.
<sup>4</sup> - There are 684 CDNs summed over all cancer types. The percentage is **n<sub>i</sub>/Total n**.

In most cancer types, 10%∼30% of patients, shown in the **n**_0_ row of **Table 4**, have no known CDNs (and >50% among breast cancer patients). Hence, the current practice is to rely on missense mutations, regardless of CDNs or non-CDNs, on the canonical genes. The CDN column vs. the gene column in **Table 4** address this issue. For example, the CDN column suggests that 33% of lung cancer patients (the **n**_0_ row) would not respond well to gene-targeted therapy whereas the gene column show only 5.3%. The difference is due to a higher, and likely inflated, detection rate of candidate drivers in the gene column. We suggest that patients who have a non-CDN mutation on a driver gene would not respond to the targeted therapy against that gene, as demonstrated in **Fig. 5**. In the above example, 27.7% (33%∼5.3%) of patients may be subjected to the targeted treatment but may not respond well.

*3) Prevalence vs. potency of CDN-bearing genes in driving tumorigenesis*

The last question is the relationship between mutation prevalence and tumorigenic strength (or potency) among CDN-bearing genes. For example, when a patient is diagnosed to have 5 CDNs in 5 genes, what may be their relative contributions to the tumorigenesis? Are they equally valid candidates for targeted therapy? It would seem logical that canonical CDGs with many CDNs should be the targets. However, because these genes would contribute at most one CDN to the tumorigenesis (**Fig. 3B**), targeting a high prevalence gene may not yield more benefits to the patients than targeting a low prevalence gene that has a CDN.

The implication is that prevalence and potency of CDNs may not be strongly correlated. Some genes may be prevalently mutated in the patient population but, in each affected patient, these genes may not be more potent than the less prevalent genes with a CDN mutation. Potency can be tested in vitro by gene editing or in vivo by targeting treatment. In this interpretation, targeting a CDN of low prevalence (say, *i* = 3) may be as effective in treatment as targeting a high prevalence CDN with *i* = 20. The model and **Table 5** present this hypothesis based on cancer hallmarks.

**Table 5.** Gene numbers for different cancer hallmarks.

| hallmark | gene number |  |  |
| --- | --- | --- | --- |
|  | all records | breast | colon |
| angiogenesis | 78 | 8 | 6 |
| cell division control | 107 | 12 | 10 |
| cell replicative immortality | 44 | 4 | 3 |
| change of cellular energetics | 70 | 10 | 4 |
| escaping immune response to cancer | 51 | 1 | 1 |
| escaping programmed cell death | 202 | 32 | 20 |
| genome instability and mutations | 106 | 10 | 7 |
| invasion and metastasis | 206 | 52 | 27 |
| proliferative signaling | 176 | 40 | 20 |
| senescence | 48 | 3 | 5 |
| suppression of growth | 130 | 11 | 12 |
| tumor promoting inflammation | 54 | 2 | 3 |

Note - data downloaded from COSMIC (https://cancer.sanger.ac.uk/cosmic/download), see methods.

The hallmarks of cancer were first proposed by (Hanahan and Weinberg 2000) with several updates (Hanahan and Weinberg 2011; Hanahan 2022). Each hallmark is a cancer phenotype shown in **Table 5** that lists the number of genes involved in each particular hallmark (see **Methods**). While each hallmark may be associated with a number of genes, many genes are also involved in multiple hallmarks. As even the highly prevalent genes would usually have at most one mutation in each patient, we assume that each gene is associated with one hallmark in each patient.

Suppose that tumorigenesis requires a mutation in most (but perhaps not all) of the hallmarks, then the number of mutation combinations would be the product of all numbers in the corresponding column. For breast cancer, it would be 8 × 12 × 4 × 11 × 2 ∼ 1.7 × 10^11^. In other words, the possible mutation combinations that can drive breast cancer is over a billion. Hence, two breast cancers are unlikely to have the same set of CDGs or CDNs. In this view, the prevalence of a gene would be inversely proportional to the hallmark gene number. For example, genes of “*invasion and metastasis*” in breast cancer would have a prevalence of < 1/52. In contrast, the potency in tumorigenesis should depend on the hallmark phenotype itself, and independent of gene number for that hallmark. In this example, each gene of “*invasion and metastasis* “ may be lowly prevalent, but could also be highly potent in each patient.

In short, the prevalence and potency of CDNs may be poorly correlated. The hypothesis can be functionally tested (by gene-editing in vitro or targeting treatment in vivo) in conjunction with the data on the attraction (i.e., co-occurrences) vs repulsion (lack of co-occurrences) of CDNs.

## DISCUSSION

The companion study presents the theory that computes the limit of recurrences (*i*/**n**, *i* times in **n** patients) of reachable by neutral mutations. Above the cutoff (e.g., 3/1000), a recurrent mutation is deemed an advantageous CDN (Zhang et al. 2024). At present, the power of CDN analysis is hampered by the still small sample sizes, generally between 300 and 3000. We show that, when **n** reaches 10^5^, a mutation only has to recur 12 times to be shown as a CDN, i.e., 25 times more sensitive than 3/1000. In short, nearly all CDNs should be discovered with **n** ≥ 10^5^.

In this study, we apply the theory on existing data to characterize the discovered CDNs. Based on the TCGA data, this study concludes that each cancer patient carries only 1∼2 CDNs, whereas 6∼10 drivers are usually hypothesized to be present in each cancer genome (Hanahan and Weinberg 2011; Vogelstein et al. 2013; Campbell et al. 2020). This deficit signifies the current incomplete understanding of cancer driving potentials. Across patients of the same cancer type, about 50 to 150 CDNs have been discovered for each cancer type, representing perhaps only 10% of all possible CDNs. Given a complete set of CDNs, it should be possible to delineate the path of tumor evolution for each individual patient.

Direct functional test of CDNs would be to introduce putative cancer-driving mutations and observe the evolution of tumors. Such a task of introducing multiple mutations that are collectively needed to drive tumorigenesis has been done only recently, and only for the best-known cancer driving mutations (Ortmann et al. 2015; Takeda et al. 2015; Hodis et al. 2022). In most tumors, the correct combination of mutations needed is not known. Clearly, CDNs, with their strong tumorigenic strength, are suitable candidates.

Many CDNs in a patient may not fall on conventional CDGs, whereas these conventional CDGs may have passenger or weak mutations. Therefore, the efforts in gene-targeting therapy may well be shifted to the CDN-harboring genes. Given a complete set of CDNs, many more driver genes can be identified. Since many driver genes cannot be targeted for biological or technical reasons (Dang et al. 2017; Danesi et al. 2021; Waarts et al. 2022), a large set of CDGs will be desirable. The goal is that each cancer patient would have multiple targetable CDGs, all driven by CDNs they carry. In that case, the probability that resistance mutations eluding multiple targeting drugs should be diminished (B. Chen et al. 2022; Zhai et al. 2022; Bian et al. 2023; Lin et al. 2023; Zhu et al. 2023).

In this context, we should comment on the feasibility of targeting CDNs that may occur in either oncogenes (ONCs) or tumor suppressor genes (TSGs). It is generally accepted that ONCs drive tumorigenesis thanks to the gain-of-function (GOF) mutations whereas TSGs derive their tumorigenic powers by loss-of-function (LOF) mutations. Nevertheless, since LOF mutations are likely to be widespread on TSGs, they are less likely to recur as CDNs. The even distributions of non-sense mutations along the length of many TSGs provide such evidence. Importantly, as gene targeting aims to diminish gene functions, GOF mutations are perceived to be targetable whereas LOF mutations are not. By extension, ONCs should be targetable but TSGs are not, an assertion we address below.

The data suggest that mis-sense mutations on TSGs may often be of the GOF kind. If mis-sense mutations are far more prevalent than nonsense mutations in tumors, the mis-sense mutations cannot possibly be LOF mutations. (After all, it is not possible to lose *more* functions than nonsense mutations.) In a separate study (Deng et al. in prep.), we compare mis-sense and nonsense mutations (referred to as the escape-route analysis). For example, AAA to AAC (K to Q) is a mis-sense mutation while the same AAA codon to AAT (K to stop) is a non-sense mutation. We found many cases where the mis-sense mutations on TSGs are more prevalent (> 10X) than nonsense mutations. We interpret these mis-sense mutations to be of the GOF kind because they could not possibly “lose” more functions than the nonsense mutations Another interesting pattern may be the distributions of CDNs across different cancer types. Cancer evolution in different tissues represents parallel evolution driven by similar selection for cell proliferation but under different ecological conditions. **Fig. 4** suggests that CDNs previously identified to be cancer-specific may have pan-cancer effects. In different cancer types, the same CDNs may drive the tumorigenesis but the strength may not be sufficient to raise the data above the statistical threshold.

The CDN approach has two additional applications. First, it can be used to find CDNs in non-coding regions. Although the number of whole genome sequences at present is still insufficient for systematic CDN detection, the preliminary analysis suggests that the density of CDNs in non-coding regions is orders of magnitude lower than in coding regions. Second, CDNs can also be used in cancer screening with the advantage of efficiency as the targeted mutations are fewer. For the same reason, the false negative rate should be much lower too. Indeed, the false positive rate should be far lower than the gene-based screen which often shows a false positive rate of >50% (supplement File S1).

Cancer evolution falls within the realm of ultra-microevolution (Wu et al. 2016). The repeated evolution addresses the single most severe criticism of evolutionary studies, namely all evolutionary events have a sample size of one. Such repeated evolution offers the opportunity to uncover the full list of mutations underling complex traits that is at the heart of molecular evolution. The genetics of speciation (Wu and Ting 2004; Pan et al. 2022; Wang et al. 2022; Wu 2023) and the emergence of major viral strains (such as COVID-19) (Deng et al. 2022; Ruan et al. 2022; Cao et al. 2023; Ruan et al. 2023) are both phenomena of complex gene interactions. The two companion studies may thus unite evolutionary biology and cancer medicine.

## ACKNOWLEDGEMENTS

We wish to acknowledge the supports from the First Affiliated Hospital, the Seventh Affiliated Hospital of Sun Yat-sen University, Cancer Center of Clifford Hospital, Jinan University, Cancer Hospital Chinese Academy of Medical Sciences, Shenzhen Center, and Guangdong Academy of Medical Sciences, Guangdong Provincial People’s Hospital on the startup of the Cancer Driving Nucleotide (CDN) project. We would like to acknowledge Kunming Institute of Zoology for discussing the ideas of CDN. We thank Weiwei Zhai, Qianfei Wang, and Weini Huang for insightful comments and suggestions. We would also like to acknowledge the American Association for Cancer Research (AACR) and The Cancer Genome Atlas (TCGA) project, which have provided invaluable datasets and resources that have significantly enriched our understanding of cancer biology and improved patient outcomes. This work was supported by the National Natural Science Foundation of China (32150006, 32293193/32293190 to C.I.W., 82341092 to HJ Wen, and 32200493 to Y.R.), the National Key Research and Development Projects of the Ministry of Science and Technology of China (2021YFC2301300, 2021YFC0863400), and Guangdong Key Research and Development Program (No. 2022B1111030001).

## Methods

### Data preparation

Single-nucleotide variant (SNV) data for TCGA patients were downloaded from the GDC Data Portal (https://portal.gdc.cancer.gov/, data version 2022-02-28), with mutations identified by at least two pipelines were included in this study. Mutations exceeding a 1‰ frequency in the Genome Aggregation Database (*gnomAD*, version v2.1.1) were excluded to minimize potential false positives arising from germline variants. Patients with more than 3000 coding region point mutations were filtered out as potential hypermutator phenotypes. This filtering process yielded a final analysis set encompassing 7369 patients across 12 diverse cancer types for subsequent analysis. The calculation of ***A***_i_ and ***S***_i_ follows the same method as described in the companion paper (https://elifesciences.org/reviewed-preprints/99340).

For CDN analysis in non-cancerous tissues, mutation profiles for normal tissues were retrieved from *SomaMutDB* (Sun et al. 2022). Mutations from different samples originating from the same individual were consolidated. Donners above the age of 80 were excluded from our dataset. The mutation processing followed the same pipeline as previously described. In total, we have mutation profiles from 487 donners serving as a negative control.

The canonical lists of cancer driver genes were obtained from three distinct data sources. The CGC Tier 1 genes, encompassing genes with the highest confidence of driver status, were retrieved from the COSMIC Cancer Gene Census (https://cancer.sanger.ac.uk/census) (Sondka et al. 2018). The IntOGen driver gene list, which employs an integrated pipeline for gene discovery, was downloaded from https://www.intogen.org/download (Martínez-Jiménez et al. 2020). Bailey’s driver gene list comprises 299 cancer driver genes identified through a *PanSoftware* strategy, with further experimental validation confirming their role in driving cell lines (Bailey et al. 2018). The consistency of cancer types across all studies was manually verified using *oncotree* (https://oncotree.mskcc.org/#/home). For the analysis of driver gene overlap, only drivers from the same cancer type were compared.

The hallmark annotation of genes is downloaded from COSMIC (https://cancer.sanger.ac.uk/cosmic/download), encompassing 331 genes with annotated dysregulated biological processes. It is important to note that these hallmarks are manually annotated as part of an ongoing effort to characterize the role of genes in cancer based on literature evidence. The actual scale of hallmark genes may be substantially larger than the current version.

For gene-level selection analysis, we utilized the R package ‘*dndscv*’ to quantify selection signals for missense and nonsense mutations in a given gene (Martincorena et al. 2017). Specifically, the package calculates the Ka/Ks ratio, denoted as ‘***w***’ in the final results, for a given mutation impact (missense or nonsense). The significance of selection is presented as *q* values after Benjamini-Hochberg (BH) adjustment. Genes with ***w*** > 1 and *q* < 0.1 were identified as being significantly under positive selection.

We employ *i*\* = 3 as a cutoff for identifying Cancer Driving Nucleotides (CDNs) across various cancer types. The specific value of *i*\* is detailed in Eq. 10 of the companion paper (Zhang et al. 2024). Here, *i*\* = 3 is chosen consistently across all cancer types, taking into account the abundance of sites under positive selection given *i* = 3 in **Table 2**. Throughout our analysis, emphasis is placed on CDNs of the missense category, where missense mutations with a recurrence ≥3 are identified as CDNs. For *ΔUi*, analysis, the reference table for 75 single-step amino acid changes was obtained from (Chen, He, et al. 2019), and the *ΔUi*, for each CDN is derived by mapping the amino acid change to the reference table.

### Calculation of *A*_i_e_

We employ Eq. 9 from the companion paper to calculate the expected value for ***A***_i_ under neutrality. For a given site, the cumulative probability for recurrence *x ≤ i* - 1 could be expressed as:

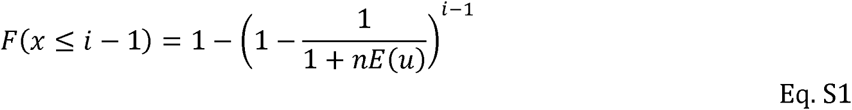

where **n** is the population size of a given cancer type, and **E(u)** is the mutation rate per site per patient derived from singleton synonymous mutations:

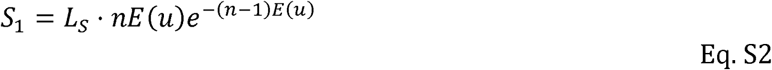

Then by expectation, site number of recurrence *i* (A*x≥i*) could be represented by:

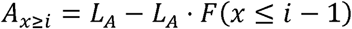

Following the same logic, we’ll have A_x≥i+1_ as:

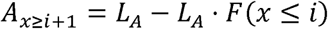

Then the expected value for ***A*_i_e_** is then:

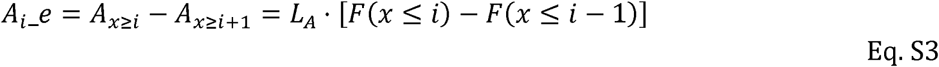

***L*** _A_ and ***L*** _S_ are missense and synonymous sites, respectively. The calculation procedure is described in methods of the companion paper (Zhang et al. 2024).

With Eqs. S1∼S3, we could solve for the expected number of sites with missense mutation recurrence *i*.

### Survival analysis of *EGFR*-targeted therapy

The mutation and clinical profiles of 23,253 patients were retrieved from the GENIE project (Cerami et al. 2012; de Bruijn et al. 2023), with 7,216 patients harboring *EGFR* mutations. Survivor months were calculated as the time elapsed between the date of sequencing and the date of the last contact (or day of death). In cases where patients had multiple sequencing reports, the earliest one was selected. For CDN calling, we applied Eq. 10 from the companion paper (Zhang et al. 2024). With ε = 0.01, we set the CDN cutoff *i*^*^ = 14. To mitigate potential biases from other common drivers in lung cancer, patients with indels in exon 19 and 20 of *EGFR*, G12/13 mutations in *KRAS*, V600 mutations in *BRAF*, exon 20 insertions in *HER2*, fusions in *MET*, *ALK*, *ROS1*, *RET*, *NTRK*, and *MET* were filtered out. The final survival analysis was conducted using *GraphPad Prism 8*.

### Annotation for non-canonical CDN genes

We conducted functional annotation and enrichment analysis for newly identified non-canonical CDN genes using four independent databases (Gene Ontology, KEGG, Disease Ontology, and Reactome) with R packages (*clusterProfiler*, *DOSE*, *ReactomePA*). For each analysis, we set a *p*-value cutoff of 0.05 and a *q*-value cutoff of 0.2, with *p* value adjustment method set to “BH”. To explore the connections between non-canonical CDN genes and canonical cancer driver genes (CDGs), enrichment analyses were performed alongside cancer drivers from IntOGen. Specifically, for enrichment annotations related to cancer hallmarks, the corresponding genes were subjected to manual confirmation using CancerGeneNET (https://signor.uniroma2.it/CancerGeneNet/).

### Data Availability

The scripts for generating the key results of this study and the accompanying paper (https://doi.org/10.7554/eLife.99340.1) are available at https://gitlab.com/ultramicroevo/cdn_v1. Example files for breast cancer analysis have also been included. The complete set of CDNs can be found in Supplementary File S2 of the accompanying paper (https://elifesciences.org/reviewed-preprints/99340).

## References

AACR Project GENIE Consortium. 2017. AACR Project GENIE: Powering Precision Medicine through an International Consortium. Cancer Discov 7:818–831.

Anandakrishnan R, Varghese RT, Kinney NA, Garner HR. 2019. Estimating the number of genetic mutations (hits) required for carcinogenesis based on the distribution of somatic mutations. PLOS Computational Biology 15:e1006881.

Armitage P, Doll R. 1954. The Age Distribution of Cancer and a Multi-stage Theory of Carcinogenesis. Br J Cancer 8:1–12.

Arnedo-Pac C, Mularoni L, Muiños F, Gonzalez-Perez A, Lopez-Bigas N. 2019. OncodriveCLUSTL: a sequence-based clustering method to identify cancer drivers. Bioinformatics 35:4788–4790.

Bailey MH, Tokheim C, Porta-Pardo E, Sengupta S, Bertrand D, Weerasinghe A, Colaprico A, Wendl MC, Kim J, Reardon B, et al. 2018. Comprehensive Characterization of Cancer Driver Genes and Mutations. Cell 173:371–385.e18.

Belikov AV. 2017. The number of key carcinogenic events can be predicted from cancer incidence. Sci Rep 7:12170.

Bian S, Wang Y, Zhou Y, Wang W, Guo L, Wen L, Fu W, Zhou X, Tang F. 2023. Integrative single-cell multiomics analyses dissect molecular signatures of intratumoral heterogeneities and differentiation states of human gastric cancer. National Science Review 10:nwad094.

Bozic I, Antal T, Ohtsuki H, Carter H, Kim D, Chen S, Karchin R, Kinzler KW, Vogelstein B, Nowak MA. 2010. Accumulation of driver and passenger mutations during tumor progression. Proceedings of the National Academy of Sciences 107:18545–18550.

de Bruijn I, Kundra R, Mastrogiacomo B, Tran TN, Sikina L, Mazor T, Li X, Ochoa A, Zhao G, Lai B, et al. 2023. Analysis and Visualization of Longitudinal Genomic and Clinical Data from the AACR Project GENIE Biopharma Collaborative in cBioPortal. Cancer Res 83:3861–3867.

Campbell PJ, Getz G, Korbel JO, Stuart JM, Jennings JL, Stein LD, Perry MD, Nahal-Bose HK, Ouellette BFF, Li CH, et al. 2020. Pan-cancer analysis of whole genomes. Nature 578:82–93.

Cao Y, Chen L, Chen H, Cun Y, Dai X, Du H, Gao F, Guo F, Guo Y, Hao P, et al. 2023. Was Wuhan the early epicenter of the COVID-19 pandemic?—A critique. National Science Review 10:nwac287.

Cerami E, Gao J, Dogrusoz U, Gross BE, Sumer SO, Aksoy BA, Jacobsen A, Byrne CJ, Heuer ML, Larsson E, et al. 2012. The cBio cancer genomics portal: an open platform for exploring multidimensional cancer genomics data. Cancer Discov 2:401–404.

Chen B, Wu X, Ruan Y, Zhang Y, Cai Q, Zapata L, Wu C-I, Lan P, Wen H. 2022. Very large hidden genetic diversity in one single tumor: evidence for tumors-in-tumor. Natl Sci Rev 9:nwac250.

Chen Q, He Z, Lan A, Shen X, Wen H, Wu C-I. 2019. Molecular Evolution in Large Steps—Codon Substitutions under Positive Selection. Molecular Biology and Evolution 36:1862–1873.

Chen Q, Lan A, Shen X, Wu C-I. 2019. Molecular Evolution in Small Steps under Prevailing Negative Selection: A Nearly Universal Rule of Codon Substitution. Genome Biology and Evolution 11:2702–2712.

Chen Q, Yang H, Feng X, Chen Qingjian, Shi S, Wu C-I, He Z. 2022. Two decades of suspect evidence for adaptive molecular evolution—negative selection confounding positive-selection signals. National Science Review 9:nwab217.

Choudhury NJ, Lavery JA, Brown S, de Bruijn I, Jee J, Tran TN, Rizvi H, Arbour KC, Whiting K, Shen R, et al. 2023. The GENIE BPC NSCLC Cohort: A Real-World Repository Integrating Standardized Clinical and Genomic Data for 1,846 Patients with Non–Small Cell Lung Cancer. Clin Cancer Res 29:3418–3428.

Danesi R, Fogli S, Indraccolo S, Del Re M, Dei Tos AP, Leoncini L, Antonuzzo L, Bonanno L, Guarneri V, Pierini A, et al. 2021. Druggable targets meet oncogenic drivers: opportunities and limitations of target-based classification of tumors and the role of Molecular Tumor Boards. ESMO Open 6:100040.

Dang CV, Reddy EP, Shokat KM, Soucek L. 2017. Drugging the “undruggable” cancer targets. Nat Rev Cancer 17:502–508.

Deng S, Xing K, He X. 2022. Mutation signatures inform the natural host of SARS-CoV-2. National Science Review 9:nwab220.

Grantham R. 1974. Amino Acid Difference Formula to Help Explain Protein Evolution. Science 185:862–864.

Hanahan D. 2022. Hallmarks of Cancer: New Dimensions. Cancer Discovery 12:31–46.

Hanahan D, Weinberg RA. 2000. The Hallmarks of Cancer. Cell 100:57–70.

Hanahan D, Weinberg RA. 2011. Hallmarks of Cancer: The Next Generation. Cell 144:646–674.

Hodis E, Triglia ET, Kwon JYH, Biancalani T, Zakka LR, Parkar S, Hütter J-C, Buffoni L, Delorey TM, Phillips D, et al. 2022. Stepwise-edited, human melanoma models reveal mutations’ effect on tumor and microenvironment. Science 376:eabi8175.

Kandoth C, McLellan MD, Vandin F, Ye K, Niu B, Lu C, Xie M, Zhang Q, McMichael JF, Wyczalkowski MA, et al. 2013. Mutational landscape and significance across 12 major cancer types. Nature 502:333–339.

Lagou V, Jiang L, Ulrich A, Zudina L, González KSG, Balkhiyarova Z, Faggian A, Maina JG, Chen S, Todorov PV, et al. 2023. GWAS of random glucose in 476,326 individuals provide insights into diabetes pathophysiology, complications and treatment stratification. Nat Genet 55:1448–1461.

Lawrence MS, Stojanov P, Polak P, Kryukov GV, Cibulskis K, Sivachenko A, Carter SL, Stewart C, Mermel CH, Roberts SA, et al. 2013. Mutational heterogeneity in cancer and the search for new cancer-associated genes. Nature 499:214–218.

Li WH, Wu CI, Luo CC. 1985. A new method for estimating synonymous and nonsynonymous rates of nucleotide substitution considering the relative likelihood of nucleotide and codon changes. Molecular Biology and Evolution 2:150–174.

Lin J, Zhan G, Liu J, Maimaitiyiming Y, Deng Z, Li B, Su K, Chen J, Sun S, Zheng W, et al. 2023. YTHDF2-mediated regulations bifurcate BHPF-induced programmed cell deaths. National Science Review 10:nwad227.

Martincorena I, Raine KM, Gerstung M, Dawson KJ, Haase K, Van Loo P, Davies H, Stratton MR, Campbell PJ. 2017. Universal Patterns of Selection in Cancer and Somatic Tissues. Cell 171:1029–1041.e21.

Martínez-Jiménez F, Muiños F, Sentís I, Deu-Pons J, Reyes-Salazar I, Arnedo-Pac C, Mularoni L, Pich O, Bonet J, Kranas H, et al. 2020. A compendium of mutational cancer driver genes. Nat Rev Cancer 20:555–572.

Meyer D, Kames J, Bar H, Komar AA, Alexaki A, Ibla J, Hunt RC, Santana-Quintero LV, Golikov A, DiCuccio M, et al. 2021. Distinct signatures of codon and codon pair usage in 32 primary tumor types in the novel database CancerCoCoPUTs for cancer-specific codon usage. Genome Med 13:122.

Mularoni L, Sabarinathan R, Deu-Pons J, Gonzalez-Perez A, López-Bigas N. 2016. OncodriveFML: a general framework to identify coding and non-coding regions with cancer driver mutations. Genome Biology 17:128.

Nei M, Gojobori T. 1986. Simple methods for estimating the numbers of synonymous and nonsynonymous nucleotide substitutions. Molecular Biology and Evolution 3:418–426.

Nik-Zainal S, Alexandrov LB, Wedge DC, Van Loo P, Greenman CD, Raine K, Jones D, Hinton J, Marshall J, Stebbings LA, et al. 2012. Mutational Processes Molding the Genomes of 21 Breast Cancers. Cell 149:979–993.

Ortmann CA, Kent DG, Nangalia J, Silber Y, Wedge DC, Grinfeld J, Baxter EJ, Massie CE, Papaemmanuil E, Menon S, et al. 2015. Effect of Mutation Order on Myeloproliferative Neoplasms. N Engl J Med 372:601–612.

Pan Y, Zhang C, Lu Y, Ning Z, Lu D, Gao Y, Zhao X, Yang Y, Guan Y, Mamatyusupu D, et al. 2022. Genomic diversity and post-admixture adaptation in the Uyghurs. National Science Review 9:nwab124.

Passaro A, Leighl N, Blackhall F, Popat S, Kerr K, Ahn MJ, Arcila ME, Arrieta O, Planchard D, De Marinis F, et al. 2022. ESMO expert consensus statements on the management of EGFR mutant non-small-cell lung cancer. Annals of Oncology 33:466–487.

Porta-Pardo E, Godzik A. 2014. e-Driver: a novel method to identify protein regions driving cancer. Bioinformatics 30:3109–3114.

Reimand J, Bader GD. 2013. Systematic analysis of somatic mutations in phosphorylation signaling predicts novel cancer drivers. Molecular Systems Biology 9:637.

Roberts SA, Gordenin DA. 2014. Hypermutation in human cancer genomes: footprints and mechanisms. Nat Rev Cancer 14:786–800.

Ruan Y, Wen H, Hou M, He Z, Lu X, Xue Y, He X, Zhang Y-P, Wu C-I. 2022. The twin-beginnings of COVID-19 in Asia and Europe—one prevails quickly. National Science Review 9:nwab223.

Ruan Y, Wen H, Hou M, Zhai W, Xu S, Lu X. 2023. On the epicenter of COVID-19 and the origin of the pandemic strain. National Science Review 10:nwac286.

Sherman MA, Yaari AU, Priebe O, Dietlein F, Loh P-R, Berger B. 2022. Genome-wide mapping of somatic mutation rates uncovers drivers of cancer. Nat Biotechnol:1–10.

Sondka Z, Bamford S, Cole CG, Ward SA, Dunham I, Forbes SA. 2018. The COSMIC Cancer Gene Census: describing genetic dysfunction across all human cancers. Nat Rev Cancer 18:696–705.

Sun S, Wang Y, Maslov AY, Dong X, Vijg J. 2022. SomaMutDB: a database of somatic mutations in normal human tissues. Nucleic Acids Research 50:D1100–D1108.

Suzuki K, Hatzikotoulas K, Southam L, Taylor HJ, Yin X, Lorenz KM, Mandla R, Huerta-Chagoya A, Melloni GEM, Kanoni S, et al. 2024. Genetic drivers of heterogeneity in type 2 diabetes pathophysiology. Nature 627:347–357.

Takeda H, Wei Z, Koso H, Rust AG, Yew CCK, Mann MB, Ward JM, Adams DJ, Copeland NG, Jenkins NA. 2015. Transposon mutagenesis identifies genes and evolutionary forces driving gastrointestinal tract tumor progression. Nat Genet 47:142–150.

Tang H, Wyckoff GJ, Lu J, Wu C-I. 2004. A universal evolutionary index for amino acid changes. Mol Biol Evol 21:1548–1556.

Tate JG, Bamford S, Jubb HC, Sondka Z, Beare DM, Bindal N, Boutselakis H, Cole CG, Creatore C, Dawson E, et al. 2019. COSMIC: the Catalogue Of Somatic Mutations In Cancer. Nucleic Acids Research 47:D941–D947.

Vogelstein B, Papadopoulos N, Velculescu VE, Zhou S, Diaz LA, Kinzler KW. 2013. Cancer Genome Landscapes. Science 339:1546–1558.

Vujkovic M, Keaton JM, Lynch JA, Miller DR, Zhou J, Tcheandjieu C, Huffman JE, Assimes TL, Lorenz K, Zhu X, et al. 2020. Discovery of 318 new risk loci for type 2 diabetes and related vascular outcomes among 1.4 million participants in a multi-ancestry meta-analysis. Nat Genet 52:680–691.

Waarts MR, Stonestrom AJ, Park YC, Levine RL. 2022. Targeting mutations in cancer. J Clin Invest 132:e154943.

Wang X, He Z, Guo Z, Yang M, Xu S, Chen Q, Shao S, Li S, Zhong C, Duke NC, et al. 2022. Extensive gene flow in secondary sympatry after allopatric speciation. National Science Review 9:nwac280.

Weinstein JN, Collisson EA, Mills GB, Shaw KRM, Ozenberger BA, Ellrott K, Shmulevich I, Sander C, Stuart JM. 2013. The Cancer Genome Atlas Pan-Cancer analysis project. Nat Genet 45:1113–1120.

Wu C-I ed. 2022. What are species and how are they formed? National Science Review 9:nwad017.

Wu C-I. 2023. The genetics of race differentiation—should it be studied? National Science Review 10:nwad068.

Wu C-I, Ting C-T. 2004. Genes and speciation. Nat Rev Genet 5:114–122.

Wu C-I, Wang H-Y, Ling S, Lu X. 2016. The Ecology and Evolution of Cancer: The Ultra-Microevolutionary Process. Annu. Rev. Genet. 50:347–369.

Xue D, Narisu N, Taylor DL, Zhang M, Grenko C, Taylor HJ, Yan T, Tang X, Sinha N, Zhu J, et al. 2023. Functional interrogation of twenty type 2 diabetes-associated genes using isogenic human embryonic stem cell-derived β-like cells. Cell Metabolism 35:1897–1914.e11.

Yang Z, Ro S, Rannala B. 2003. Likelihood Models of Somatic Mutation and Codon Substitution in Cancer Genes. Genetics 165:695–705.

Yang Z, Swanson WJ. 2002. Codon-Substitution Models to Detect Adaptive Evolution that Account for Heterogeneous Selective Pressures Among Site Classes. Molecular Biology and Evolution 19:49–57.

Zhai W, Lai H, Kaya NA, Chen J, Yang H, Lu B, Lim JQ, Ma S, Chew SC, Chua KP, et al. 2022. Dynamic phenotypic heterogeneity and the evolution of multiple RNA subtypes in hepatocellular carcinoma: the PLANET study. National Science Review 9:nwab192.

Zhang L, Deng T, Liufu Z, Liu X, Chen B, Hu Z, Liu C, Lu X, Wen H, Wu C-I. 2024. The theory of massively repeated evolution and full identifications of Cancer Driving Nucleotides (CDNs). eLife [Internet] 13. Available from: https://elifesciences.org/reviewed-preprints/99340

Zhu H, Lin Y, Lu D, Wang S, Liu Y, Dong L, Meng Q, Gao J, Wang Y, Song N, et al. 2023. Proteomics of adjacent-to-tumor samples uncovers clinically relevant biological events in hepatocellular carcinoma. National Science Review 10:nwad167.

