## supplement file S1 for "On the discovered Cancer Driving Nucleotides (CDNs)–Distributions across genes, cancer types and patients"

**1. Quantifying evolutionary fitness of CDN.**

We leverage Eq. 2 and Eq. 4 from the companion paper (Zhang et al. 2024) and rewrite ***A_i_*** as follows:

$$A_{i}=A_{i}^{neut}+A_{i}^{*}$$

Eq. S1

Where ***A_i_*** represents the observed site number with missense recurrence of *i*, which could be further decomposed into 2 components: $A_{i}^{neut}$: the site number with missense recurrence of *i* under neutral mutational force, and $A_{i}^{*}$, which occurs under positive selection. For $A_{i}^{*}$, we have:

$$A_{i}^{*}=f\cdot L_{A}\cdot g\left( i,k \right)\left[ w\cdot nE\left( u \right) \right]^{i}$$

Eq. S2

With Eq. S1 and S2, ***A_i_*** could be expressed as:

$$A_{i}\sim\frac{\Gamma\left( k+i \right)}{\Gamma\left( i+1 \right)\Gamma\left( k \right)}\cdot\frac{L_{A}}{k^{i}}\left[ nE\left( u \right) \right]^{i}\cdot\left( 1+f\cdot w^{i} \right)=2.3\cdot S_{i}\cdot\left( 1+f\cdot w^{i} \right)$$

Eq. S3

Where *f* denotes the fraction of missense sites under positive selection (*f* ≪ 1), and ***w*** represents the selective advantage (*s*) scaled by the population size of progenitor cancer cell (*N*). First, we aimed to estimate *f* from the discrepancy between $R_{0}=A_{0}/S_{0}$ and $R_{1}=A_{1}/S_{1}$. The number of sites under positive selection in ***A_1_*** ($\boldsymbol{A}_{\boldsymbol{1}}^{\boldsymbol{*}}$) could be approximated from ***A_0_*** as $A_{0}\cdot f$, which could also be derived from the excess of mutations from ***A_1_*** as $A_{1}\cdot\left( R_{1}-R_{0} \right)/R_{1}$, then we have:

$$A_{0}\cdot f=A_{1}\cdot\left( R_{1}-R_{0} \right)/R_{1}$$

$$f=\frac{A_{1}}{A_{0}}\cdot\left( 1-\frac{R_{0}}{R_{1}} \right)$$

Eq. S3

Based on the average statistics in **Table 1**, *f* could be estimated from Eq. S3 to be 3.13 × 10^-4^. With synonymous recurrence sites as neutral reference, $A_{i}\sim2.3\cdot S_{i}\cdot\left( 1+f\cdot w^{i} \right)$. Given ***A_3_*** / ***S_3_*** = 5.25, we’ll have ***w*** = 16, which means ***A_3_*** would be 4096 (16^3^) fold higher than the neutral expectation.

**2. Functional annotation of new cancer drivers.**

A limitation in cancer driver discovery lies in the modelling of background mutation process, which often necessitates a balance between current knowledge and unknown mutational mechanisms. Consequently, genes recognized as canonical drivers in one cancer type may be categorized as non-canonical in others due to the lack of statistical significance. Of the 229 non-canonical drivers identified across six cancer types, 19 genes have been previously recognized as canonical drivers in different cancer types, while 23 genes were classified as drivers in IntOGen through a combination of diverse statistical methods.

For the newly identified non-canonical CDN genes in this study, we explore their potential functional relevance to cancer through a two-step procedure. First, we annotate these genes considering gene ontology, pathway, disease association and protein-protein interaction with known cancer drivers. Subsequently, we conduct manual curation by reviewing published literatures for evidence related to cancer. **Fig. S1** illustrates how non-canonical CDGs are enriched in cancer-related biological processes in lung and colon cancers. The results reveal that processes such as cell migration / adhesion, epithelial-to-mesenchymal transition (EMT), cell proliferation, energy metabolism, immune response, and DNA transcription activity are among the most enriched processes in both cancer types. Additionally, other cancer hallmark-related processes, such as the cell cycle control, DNA stability, and response to stresses, are also enriched among non-canonical CDN genes.

In **Fig. S2**, we present the top 10 genes being most connected to known cancer drivers across four independent enrichment analyses. Here, we take two unidentified driver genes for example to illustrate their functional roles relating to cancer. *PIK3R2* (Phosphoinositide-3-Kinase Regulatory Subunit 2), which encodes p85β of class I PI3K, is often highly expressed in most tumors (Liu et al. 2022). This gene has a CDN mutation of G1117A (with amino acid change of G373R) in endometrium cancer, which is also presented in lung and urinary tract cancers. *PIK3R2* has been reported as an oncogene, with its overexpression triggering cell transformation in culture and promoting cancer progression in mouse model (Vallejo-Díaz et al. 2019). *SLC7A5* (Solute Carrier Family 7 Member 5), with a CDN mutation of G1480A (V494I) in colon cancer, plays a critical oncogenic role in maintaining intracellular amino acid levels for an elevated protein synthesis rate in *KRAS*-mutant cells. Depletion of *SLC7A5* suppresses intestinal tumorigenesis in mice and resensitizes tumors to protein synthesis inhibition (Najumudeen et al. 2021). In conclusion, although excluded from the canonical driver list due to the lack of statistical significance, non-canonical CDN genes may still undergo positive selection at the site level. Ongoing research may provide further experimental evidence for these genes as part of an ongoing effort to identify the complete set of cancer drivers.

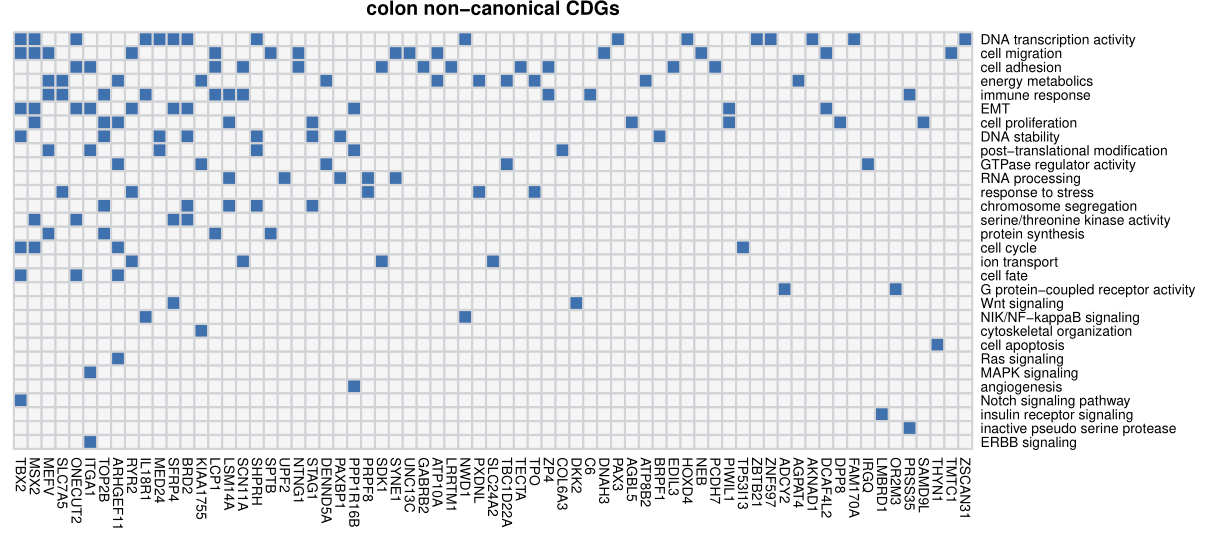

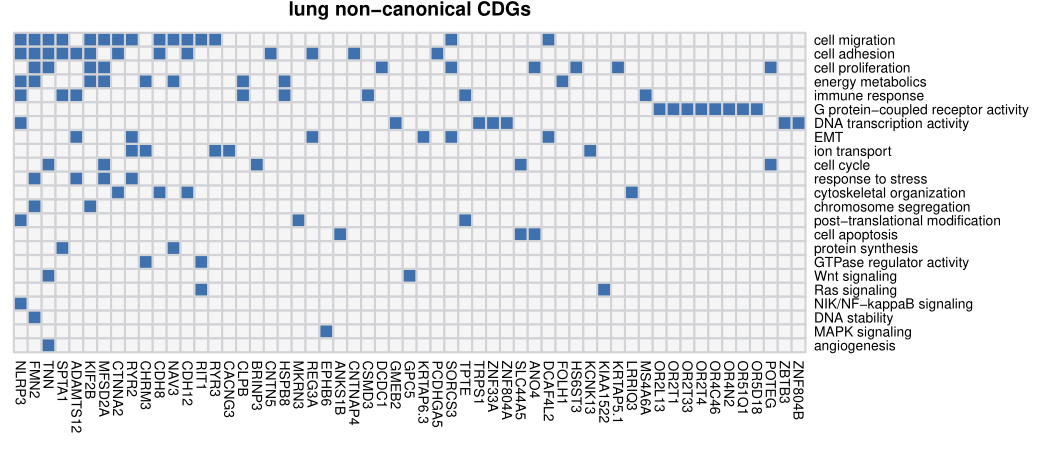

**Figure S1**: Non-canonical CDGs in colon and lung cancer along with associated biological processes (Y-axis). For each gene, we examine its annotation results from GO analysis and search for cancer-related evidence in the literature. Biological processes are summarized and curated in relation to cancer hallmarks. Each connection between gene ID and biological process is depicted by a blue block in the grid.

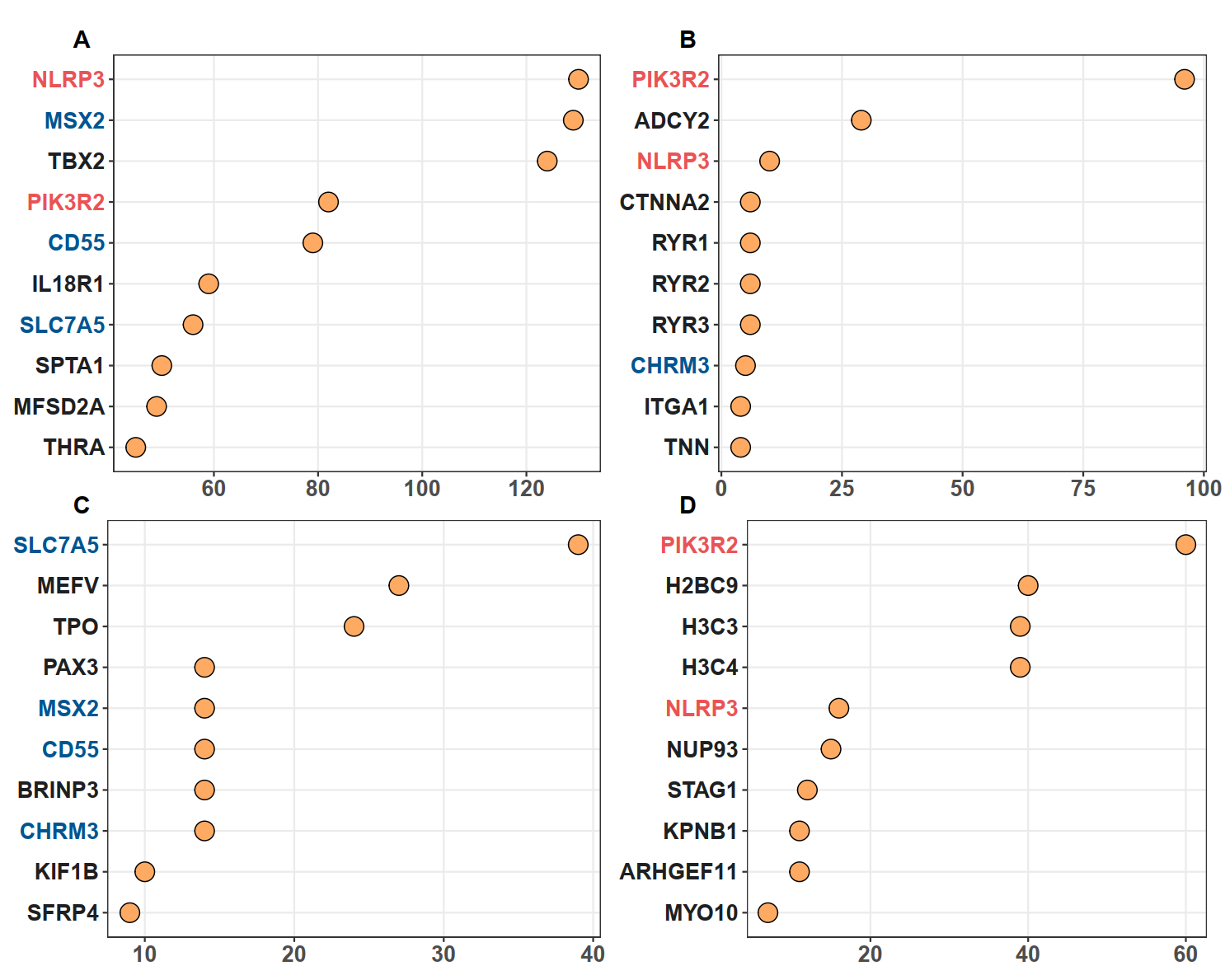

**Figure S2**: Top 10 non-canonical CDGs with the highest enrichment records with IntOGen's driver list from four enrichment analysis. Panels (**A-D**) corresponds to Gene Ontology, KEGG, Disease Ontology and Reactome analysis, respectively. The X-axis represents the number of enrichment records for each gene, while genes are listed on the Y-axis according to their enrichment record number. Genes with different occurrences across the top set of four analysis are marked with red (3 hits), blue (2 hits) and black (1 hit).

**3. The specificity of CDNs in cancer detection.**

Generally, in a given sample, each mutated gene would harbor one mutation. Therefore, we measure the false positive rate for CDNs (or CDGs) as the proportion of individuals harboring nonsynonymous mutations at CDN (or CDGs). Across 487 individuals from non-cancerous set, 446 are devoid of any mutations at CDNs, yielding a specificity of 91.6% (false positive rate: 8.4%). In contrast, for CDGs downloaded from CGC, the specificity is 37.6% (false positive rate: 62.4%). The high specificity implies the potential application of CDNs in biopsy and companion diagnostics, as well as the possibility of integration with other early screening pipelines. Furthermore, when we compared the mutations of recurrences *i* ≥ 3 in *SomaMutDB* with CDNs identified in our analysis, no overlap was observed. The high exclusiveness of CDNs between cancer and non-cancer implies that positive selection operates in a specific manner in cancer, distinct from normal tissues.

**4. the overlapping between cancer driver gene lists.**

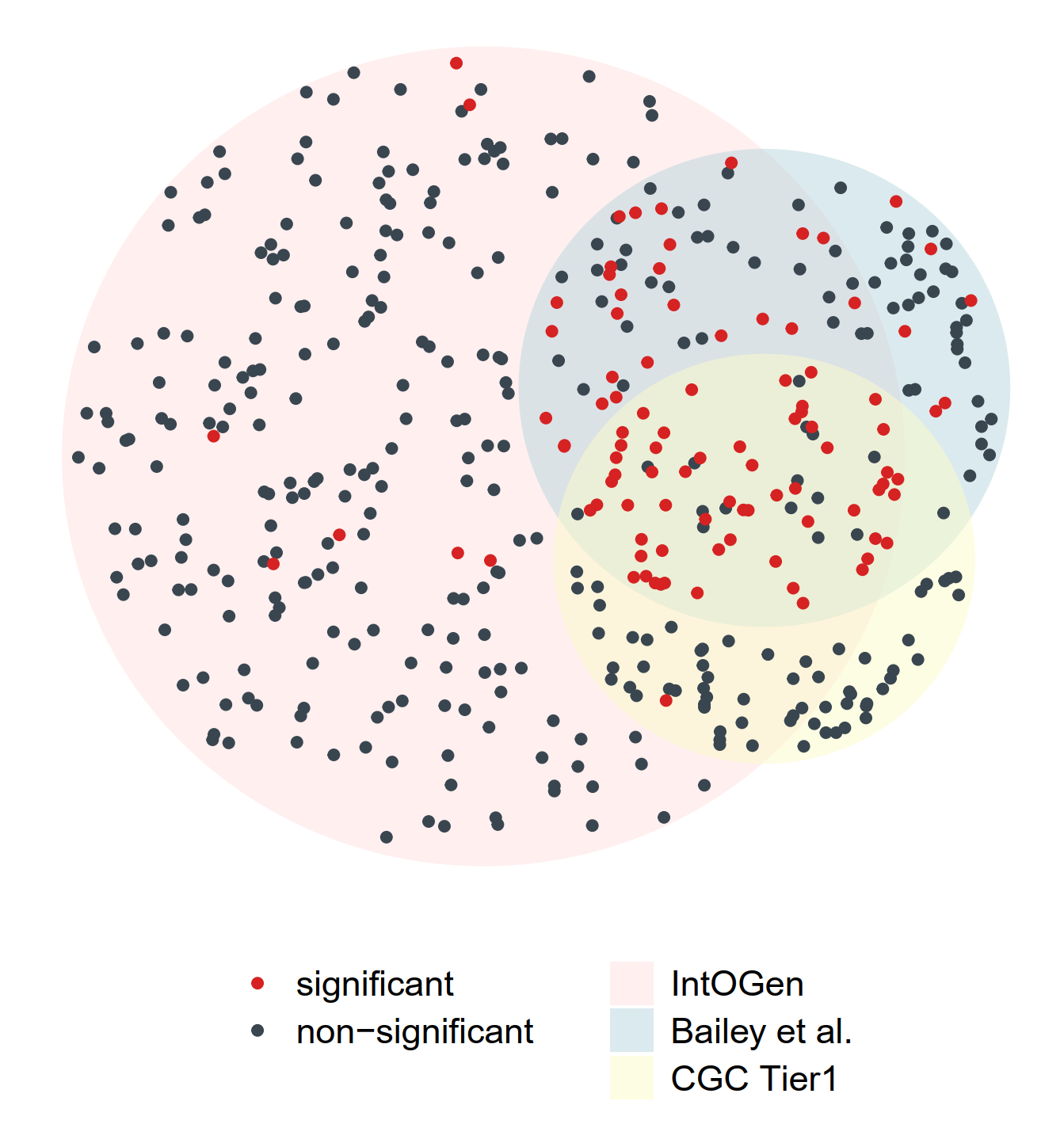

**Figure S3**: The overlap of cancer drivers from IntOGen, Bailey et al. and CGC Tier1(Bailey et al. 2018; Sondka et al. 2018; Martínez-Jiménez et al. 2020). Driver genes (dots) for 12 cancer types were extracted from each driver list, indicated by three different region colors. The area size of each region is proportional to the gene number, with 384 genes for IntOGen, 168 for Bailey et al., and 137 for CGC Tier1. Genes with a significant positive selection signal in the merged mutation set are marked in red, while non-significant ones are colored in blue. Notably, genes shared across the three driver sets are largely those with a significant Ka/Ks > 1.

**References**

Bailey MH, Tokheim C, Porta-Pardo E, Sengupta S, Bertrand D, Weerasinghe A, Colaprico A, Wendl MC, Kim J, Reardon B, et al. 2018. Comprehensive Characterization of Cancer Driver Genes and Mutations. *Cell* 173:371-385.e18.

Liu Y, Wang D, Li Z, Li X, Jin M, Jia N, Cui X, Hu G, Tang T, Yu Q. 2022. Pan-cancer analysis on the role of PIK3R1 and PIK3R2 in human tumors. *Sci Rep* 12:5924.

Martínez-Jiménez F, Muiños F, Sentís I, Deu-Pons J, Reyes-Salazar I, Arnedo-Pac C, Mularoni L, Pich O, Bonet J, Kranas H, et al. 2020. A compendium of mutational cancer driver genes. *Nat Rev Cancer* 20:555–572.

Najumudeen AK, Ceteci F, Fey SK, Hamm G, Steven RT, Hall H, Nikula CJ, Dexter A, Murta T, Race AM, et al. 2021. The amino acid transporter SLC7A5 is required for efficient growth of KRAS-mutant colorectal cancer. *Nat Genet* 53:16–26.

Sondka Z, Bamford S, Cole CG, Ward SA, Dunham I, Forbes SA. 2018. The COSMIC Cancer Gene Census: describing genetic dysfunction across all human cancers. *Nat Rev Cancer* 18:696–705.

Vallejo-Díaz J, Chagoyen M, Olazabal-Morán M, González-García A, Carrera AC. 2019. The Opposing Roles of PIK3R1/p85α and PIK3R2/p85β in Cancer. *Trends in Cancer* 5:233–244.

Zhang L, Deng T, Liufu Z, Liu X, Chen B, Hu Z, Liu C, Lu X, Wen H, Wu C-I. 2024. The theory of massively repeated evolution and full identifications of Cancer Driving Nucleotides (CDNs). *eLife* [Internet] 13. Available from: https://elifesciences.org/reviewed-preprints/99340
